# Ancestral splice variation is a key substrate for rapid diversification in African cichlids

**DOI:** 10.1101/2025.05.13.653813

**Authors:** Pooja Singh, Ehsan P. Ahi, Anna Duenser, Marija Durdevic, Wolfgang Gessl, Sylvia Schaeffer, Julian Gallaun, Ole Seehausen, Christian Sturmbauer

## Abstract

Adaptive radiation is a major driver of biodiversity. In some of the largest radiations, the pace of morphological change far outstrips the protein-coding mutation rate, suggesting that gene regulation may be important for diversification. Yet, the specific gene regulatory mechanisms shaping adaptive radiation remain poorly understood. Analysing of 200 transcriptomes from three independent but phylogenetically nested African cichlid adaptive radiations we show that alternative splicing (AS) evolved faster than gene expression (GE), playing a pivotal role in shaping novel trophic adaptations in the youngest and most species rich radiations in Lakes Victoria and Malawi. This divergence was largely driven by ancestral alternative isoforms, which, though present at low levels in related lineages that did not form radiations, increased in frequency during adaptive radiation. In addition, novel isoforms of craniofacial genes - some evolving within just a few thousand years contributed further to adaptation. The rapid turnover of AS is consistent with periods of relaxed selection followed by directional selection on alternative isoforms and splice sites, a dynamic that may have preserved a rich cache of isoform variation in these radiations and enabled ecological diversification as adaptive zones became available. We argue that the interplay between splicing and different forms of selection facilitates the generation and maintenance of protein-coding diversity, promoting evolutionary innovation into many ecologically different species at extremely short timescales.

**Significance statement:** Adaptive radiation—when species rapidly diversify to fill new ecological niches—is a key driver of biodiversity but its underlying molecular mechanisms remain unclear. Using comparative transcriptome sequencing across multiple African cichlid fish adaptive radiations, we found that rapid changes in alternative splicing—the process generating different protein-coding isoforms from the same gene—contributed more to early ecological divergence than shifts in gene expression levels. Most adaptive isoforms were present at low levels in the ancestor of these fishes but some new isoforms evolved remarkably fast, shaping diverse ecologies. We show that alternative splice variation, often thought to be biological "noise," can be a powerful and labile source of rapid evolutionary innovation during early stages of adaptive radiation.

## Introduction

Adaptive radiations are characterised by bursts of ecological and morphological diversity that have shaped the biodiversity on our planet (1). Rates of molecular evolution in protein-coding genes cannot explain the rate at which morphological evolution happens during adaptive radiation (2–4), which is why gene regulation is hypothesised to be an important mechanism for ecomorphological change in these lineages (5–7). Transcriptional changes in gene expression (GE) fine-tune the amount of messenger RNA (mRNA) and thus proteins produced from a gene (3, 8) and alternative splicing (AS) is a co-/post-transcriptional mechanism that can rapidly diversify the kinds of transcript and protein isoforms coded by a single gene by generating multiple distinct messenger RNAs (mRNAs) (9, 10). While some inroads have been made (7, 11–15), we still understand little about how these transcriptional and post-transcriptional regulatory mechanisms interact and are shaped by evolutionary processes to give rise to species-rich bursts of adaptive radiation (5).

To address this knowledge gap, here we examined the evolutionary dynamics of GE and AS in cichlid fishes from Lake Victoria, Lake Malawi and Lake Tanganyika that represent three of the fastest and ecomorphologically most diverse vertebrate adaptive radiations. Species in these lakes have evolved diverse jaws and feeding habits to adapt to different food sources, forming species-rich communities spanning complex foodwebs (16, 17). Across lakes, similar feeding ecologies and morphologies have evolved repeatedly, suggesting that cichlid lineages solved ecological challenges in similar ways, and that cichlid evolution may be constrained (18–23). We sequenced whole transcriptomes of oral (OJ) and pharyngeal jaws (PJ), the functional decoupling of which is considered a key evolutionary innovation that opened up novel trophic niches for cichlid fishes (24), from 18 phylogenetically nested cichlid species. We selected species with ecologically divergent herbivorous and carnivorous trophic adaptations within lakes and ecologically convergent trophic adaptations across lakes (16, 19, 20) (Figure 1A). This allowed us to address dynamics of repeated evolution as well as divergent evolution. Our sampled lineages represent the evolutionary process of adaptive radiation at different stages – Lake Victoria is the youngest and has the fastest rates of speciation, Lake Malawi is middle-aged, and Lake Tanganyika is the oldest and most ecomorphologically and genetically distinct (25, 26) – providing a powerful framework to study the temporal role of different gene regulatory mechanisms. We also included data from two non-radiating cichlid species in our study that live in these lakes, and thus has access to ecological opportunity, but did not form radiations. These non-radiating *Astatotilapia*/*Astatoreochromis* lineages are also phylogenetic sister-groups to the radiations, with whom they share a recent common ancestor (25, 27). All our selected species are from the tribe Haplochromini, which is the most species-rich and ecologically diverse cichlid group (28) (Figure 1B for phylogenetic relationships). The ‘Out of Tanganyika’ theory of East African cichlid evolution suggests that an ancient riverine *Astatoreochromis*-like haplochromine lineage from Lake Tanganyika, colonised Lake Malawi and Lake Victoria, contributing to those radiations, and then re-entered Lake Tanganyika to give rise to the haplochromine radiation of Tropheini cichlids (27). It has been hypothesised that ancestral standing variation may have been an important driver of adaptive radiation in East African cichlids (29–31). Based on this theory, and through the sampling of lineages that formed radiations versus those that did not, we tested whether ancestral variation in GE and AS (1) contributed to divergent and convergent trophic evolution during adaptive radiation, (2) play distinct roles in biological diversification, such as rapid functional diversfication during/near speciation through splicing versus much slower adaptive fine-tuning after speciation through gene expression, and finally (3) if we can find evidence for directional selection on altenative isoforms of craniofacial genes.

**Figure 1.**
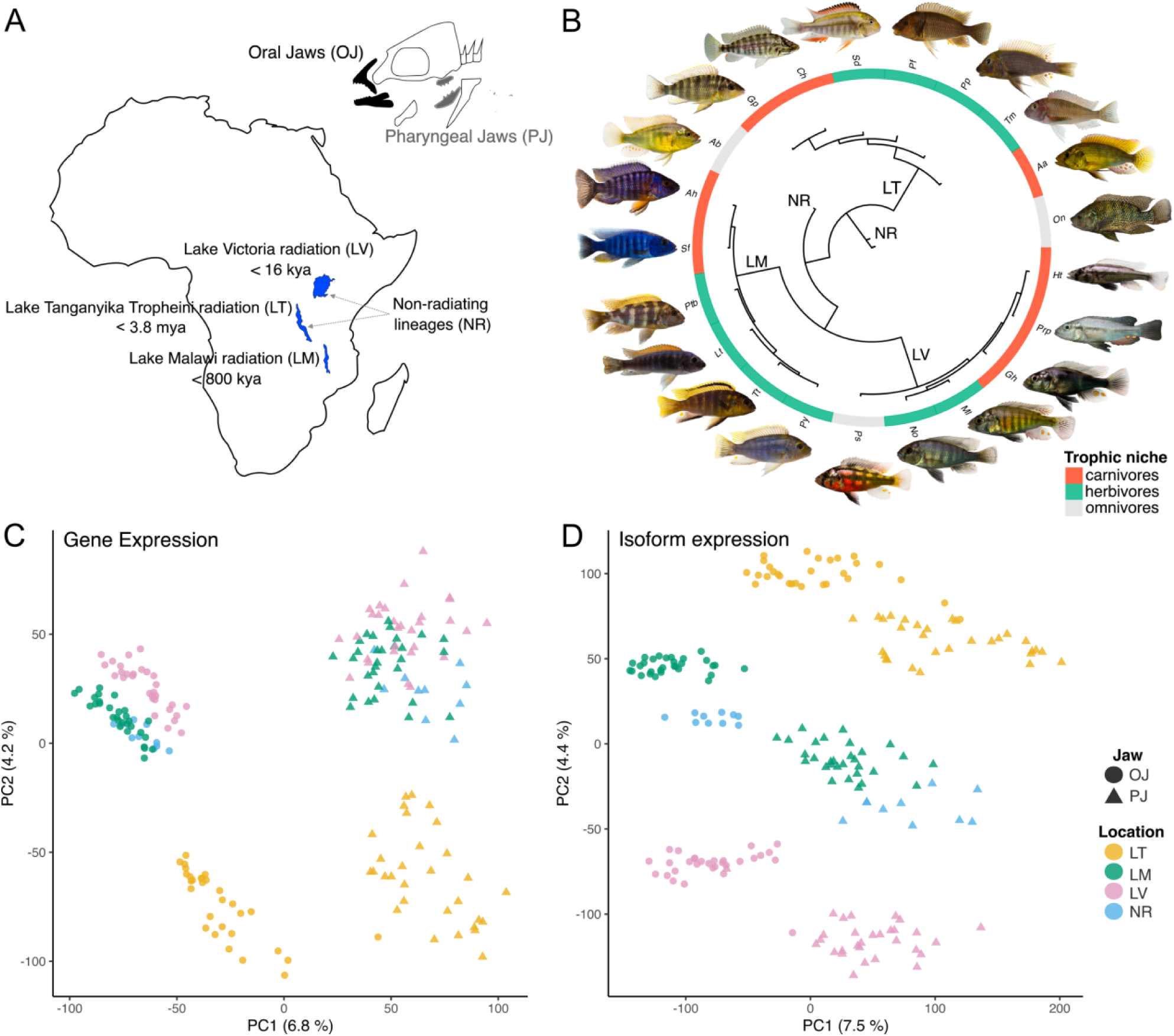
Study overview. (**A, B)** Biogeographical setting of haplochromine cichlid species sampled in this study (20 species, 2 jaw tissues (oral and pharyngeal), 5 biological replicates per species per tissue, n=200). Phylogenetic relationships reconstructed using transcriptome SNPs. Lake Victoria: *Paralabidochromis sauvagei (*Ps), *Neochromis omnicaeruleus* (No), *Mbipia lutea* (Ml), *Gaurochromis hiatus* (Gh) *Prognathochromis perrieri (*Prp); *Haplochromis thereuterion* (Ht); Lake Malawi: *Aulonocara hansbaenschi* (Ah), *Sciaenochromis fryeri* (Sf), *Petrotilapia* sp. *’Thick bars’ (*Ptb), *Labeotropheus trewavasae* (Lt), *Tropheops tropheops* (Tt), *Petrotilapia* sp. ‘yellow-chin’ (Py); Lake Tanganyika: *Tropheus moorii* (Tm), *Petrochromis polyodon* (Pp)*, Petrochromis famula* (Pf), *Simochromis diagramma* (Sd), *Ctenochromis horei* (Ch), *Gnathochromis pfefferi* (Gp). Non-radiating (NR) haplochromines, *Astatotilapia burtoni* (Ab) and *Astatoreochromis alluaudi* (Aa) are included as riverine lineages that entered lakes but did not radiate, and are also phylogenetic sister lineage to the radiations. *O. niloticus* (On) is the outgroup species which was included for phylogenetic analyses only. (**C, D)** Principal component analysis of transcriptome-wide normalised gene and isoform expression from oral (OJ) and pharyngeal jaws (PJ) of all 20 species.

## Results

### Global patterns of gene and isoform expression

To study the evolution of GE and AS during adaptive radiation, we generated transcriptomes of two tissues (OJ and PJ) from 100 cichlid fishes belonging to 20 species (Figure 1, Table S1) from three lake adaptive radiations and two species that entered lakes but did not make radiations (5 individuals per tissue per species, total N = 200). Reads were mapped to the phylogenetically equidistant *Oreochromis niloticus* reference genome(32) (median sequencing depth per tissue: median ∼6.4 million 125 base-pair (bp) paired end reads per library; median ∼80% uniquely mapped read; Table S1) using the genome-guided transcript assembly function. Gene annotations of species and tissues constructed using StringTie(33) were merged incrementally into one super-annotation(33) that was used for downstream analysis. This strategy helped to reduce false positive while increasing detection of novel and lowly expressed isoforms. Gene and isoform expression was quantified using established methods(33, 34).

To explore global patterns of gene and isoform expression, we performed Principal Component Analysis (PCA) across all samples for all genes. Gene expression PC1 separated the two tissues, and on PC2, Lake Tanganyika clustered separately from the younger radiations from Lakes Malawi and Victoria and the non-radiating species, which formed an overlapping cluster (Figure 1C). PCA clustering of isoform expression, which is a consequence of AS, was also tissue-specific on PC1, but clearly lake-specific on PC2 (Figure 1D) with non-radiating species clustering closer to Lake Malawi samples, particularly for the PJ. Both gene and isoform expression patterns within tissues captures structure among radiations (Figure 1B), with stronger signals emerging from the isoform expression. These patterns suggest that isoform-level variation may capture interspecific differences, even among very young species-flocks, that are not apparent at the gene expression level, although PCA provides only qualitative indication of these relationships.

### Splicing and expression have contrasting evolutionary dynamics

Alternative splicing results in the change of relative abundance of isoforms produced from the same gene(9), which may have evolutionary consequences(5, 35). But little is known about how AS evolves at difference timescales during adaptive radiation. To adress this, we calculated percent spliced in (PSI), a standard AS metric that represents the relative abundance of an isoform as a proportion of the total GE(36, 37) and computed Spearman’s rank correlation coefficients (Spearman’s ρ) across all samples and tissues for GE (all genes) and PSI (only genes with more than one isoform) to compare rate of change among these two processes. Our analysis revealed higher Spearman’s ρ values for GE than AS (Figure 2 boxplot, two-sided t-test *p* < 0.001). This suggets that GE was more conserved (less divergent) across samples than PSI during cichlid evolution; while AS was evolving faster. We next tested if constitutively expressed isoforms were more conserved during cichlid evolution than alternative isoforms. Indeed, we found that Spearmans’s ρ of PSI for alternative isoforms was significantly lower than that of constitutive isoforms (Figure S1, two-sided t-test *p* < 0.001). This fits the expectation that transcriptome-wide alternative isoforms are evolving under relaxed selection pressures during adaptive radiation ((38, 39).

**Figure 2.**
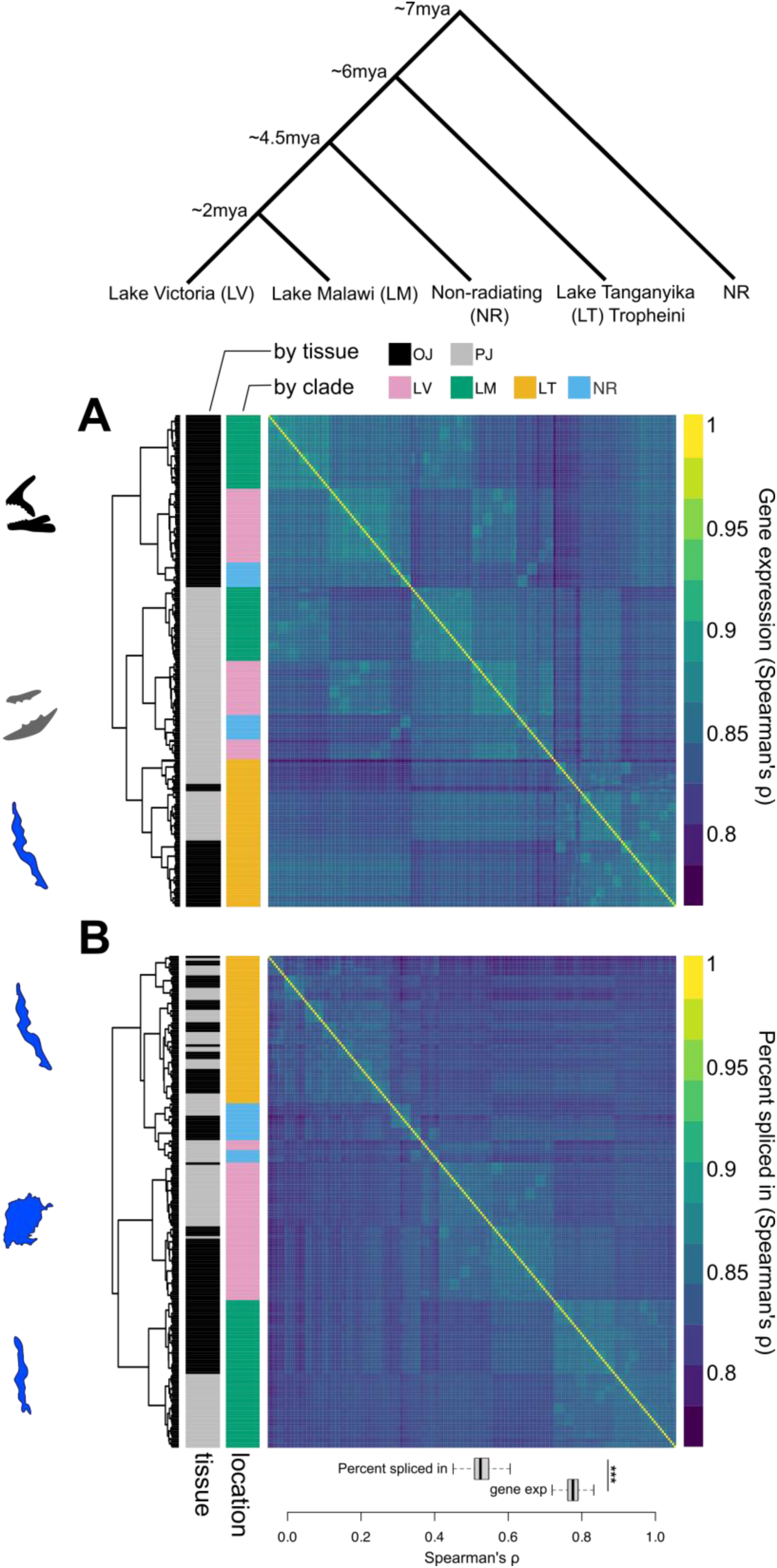
Profiling of gene expression and alternative splicing in East African cichlids. Hierarchical clustering of pairwise Spearman’s rank correlation coefficient (ρ) of transcriptome-wide (**A)** normalised gene expression (GE) and (**B)** isoform percent spliced in (PSI) of genes with more than one isoform. Analysis was conducted across all samples (n=200). Sample clustering is represented as a tree on the left side of the heatmaps, with tissue and clade (radiations or nonradiating) of origin of species annotated in colours and the dominant cluster signature illustrated as radiation/non-radiation/jaw. Boxplots show significant difference in distribution of Spearman’s ρ of PSI and GE across all samples (two-sides t-test *p* < 0.001). See Figure S1A for boxplots of Spearman’s ρ of PSI and GE calculated per lake. Cladogram illustrates simplified cladogram of phylogenetic relationship of species shown in Figure 1B. Divergence times taken from Irisarri*, Singh* et al (2018) ((25). See Figure S2 for sample names.

Next, we analysed the hierarchical clustering of Spearman’s ρ for GE and PSI. We found that GE grouped samples by tissue for younger radiations (Lakes Victoria and Malawi) and nonradiating species (Ab, Aa). However, for the older Lake Tanganyika Tropheini radiation, samples clustered first by clade/radiation and then by tissue within the radiation (Figure 2A), defying well-established expectations of tissue-specific conservation of GE in vertebrates(40). In contrast, Spearman’s ρ of PSI clustered samples first by clade/radiation, regardless of tissue, and samples from Lake Tanganyika and nonradiating species further clustered by species, regardless of tissue (Figure 2B, S2). The latter pattern would be expected if changes in splicing were evolutionarily conserved (heritable) in species and clades. This suggests that clade-specific divergence in splicing exceeds tissue-specific conserved signatures of splicing at a phylogenetic distance of ∼2mya, and species-specific divergence in splicing exceeds tissue-specific divergence at ∼6mya. Hierachical clustering relationships generally reflected phylogenetic relationships of the radiations, however, non-radiating species clustered with Lake Victoria in the gene expresssion analysis but with Lake Tangayika in the PSI analysis. Interestingly, PJ samples of *H. thereuterion* from Lake Victoria were placed outside of the Lake Victoria clade, sister to the PJ’s of the non-radiating *A. burtoni*, which is distributed in Lake Tanganyika basin and not found in Lake Victoria. Taken together with the PCA results (Figure 1A,B), our results indicate that AS may have played a more important role in the evolution of clade and species-specific jaw specialisations compared to GE. It may further suggest that heritable changes in splicing emerge rapidly during/near speciation in cichlid adaptive radiations, while GE changes may fine-tune morphology over longer evolutionary timescales.

### Genes and isoforms evolve under stabilising selection

Next, we investigated the evolution of gene and isoform expression along the phylogeny (constructed using SNPs from the RNAseq data, Figure S3A) by fitting Brownian motion (BM) and Ornstein-Uhlenbeck (OU) processes. Ornstein-Uhlenbeck has been applied to model stochastic GE evolution towards a single optimum i.e. stabilising selection(41). Brownian motion models neutral evolution i.e. genetic drift. The distribution of OU variance across the phylogeny, denoted ‘evolutionary variance’(42), depicts how constrained expression evolution is. We found that expression of most genes and isoforms was evolving under stabilising selection (90–92%) and 8-10% of genes were evolving neutrally (chi-squared test *q*-value < 0.05, Figure S3B, File S1). This has been previously reported for gene expression model species(41) but not for isoform expression. However, in contrast to gene expression, isoform expression had significantly higher evolutionary variance (less constraint) than GE under the OU model for both jaws (Figure S3C). This higher variance could be the result of weaker stabilising selection (or strong relaxed selection or perhaps the result of complex mix of selection) acting on alternative isoform expression than constitutive isoform expression, the latter being under stabilising selection to preserve function. These results reinforce the idea of fast evolving AS during adaptive radiation.

### Most splice variation is ancestral, but some may be novel

Ancestral standing variation is proposed to fuel rapid cichlid adaptive radiations(31). One way that this could happen is through the sorting of ancestral variation in gene regulation(43), such as splice variation. As a first step to test if ancestral splice variation contributed to trophic divergence, we quantified the number of genes and isoforms expressed uniquely in each radiation versus those shared between radiations and non-radiating species. The total number of both genes and isoforms expressed in the OJ was higher than in the PJ; and regardless of tissue, the absolute number of genes expressed were significantly higher in samples from the younger radiations (Lakes Malawi and Victoria) and the non-radiating lineages than the older radiation of Lake Tanganyika (Mann-Whitney U test *p* < 0.001; Figure 3A,B; S4 A,B). This overall pattern was also observed when we analysed the number of isoforms per gene (Figure S5) but the difference was only significant between the PJ’s of Lake Victoria and Malawi (Mann-Whitney U test *p* < 0.05). We tested the homogeneity of variance among groups using the Barlett’s test and found that significant differences for the number of isoforms per gene for PJ (*p* = 0.017) but not the OJ. This suggests that while the absolute number of isoforms expressed per gene is reduced over time, the transcriptional variance among species increases with time during adaptive radiation. To understand how much of this expression variation was ancestral versus putatively *de novo*, we investigated if each gene or isoform was expressed in only one of the clades, or across several radiating and non-radiating clades (Figure 3C). The majority of genes (86.0%) and isoforms (73.0%) were expressed in at least one non-radiating species and one or more radiations – this variation is likely ancestral to all East African haplochromines. These results are consistent with the ‘Out of Tanganyika’ hypothesis (27) – where similar pool of ancestral variation would be presumably present at the base of all these three radiations, and also in the non-radiating haplochromine lineages. However, in addition to this “Out of Tanganyika” colonisation (27), ancestral hybridisation from different divergent lineages from the Congo and Nile (that are also haplochromine cichlids) have shown to contribute genetic material independently to all of these radiations (25, 44, 45), which would certainly have impacted the transcriptional landscape within each radiations. Furthermore, demographic history of these radiations probably also plays a role, as the Lake Victoria has dried up and refilled (most recently 16,000 years ago) and this would impact the shared gene regulatory variation across all three lakes. All these factors probably contribute to why so little gene/isoform expression is shared across all three radiations, but larger sharing is observed in different subsets of radiating/non-radiating species. Given what we know about their phylogenetic relationships (Figure 1B), the most parsimonious explanation is that this variation was ancestral and was lost in some radiations; instead of being independently gained in non-radiating lineages and a subset of radiation. A small subset of isoforms (∼12.0%) were unique to one radiation only – pointing towards the evolution of novel splice variation in the course of adaptive radiation – or less likely, repeated loss in all other lineages. Novel isoforms can rapidly evolve through weak purifying selection on alternative splice sites ((39). It is of course possible that these putatively novel isoforms may be ancestral but the pattern is missed by our sample selection. It is also possible that isoform variation arrived through hybridisation, as cycles of hybridisation have fueled cichlid adaptive radiations ((25, 26, 45, 46). Taken together, our data support the hypothesis that substantial transcriptomic variation existed in the common ancestor of all East African haplochromines that was likely shaped by selection and drift during adaptive radiation, with some isoforms being purged and others retained; with some putatively novel isoforms evolving in the course of adaptive radiation. But did these ancestral and novel isoforms contribute to ecological adaptation in cichlid radiations?

**Figure 3.**
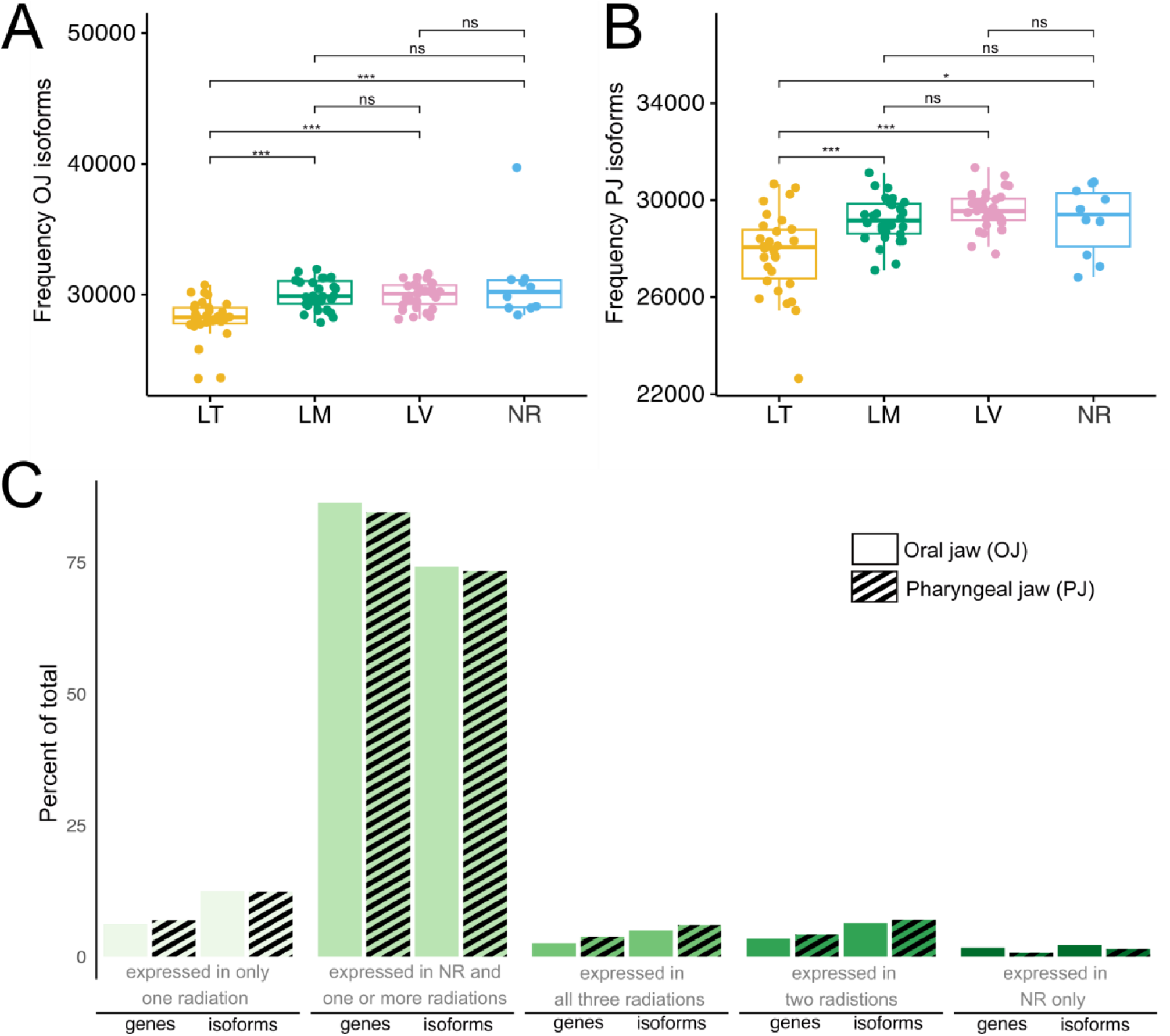
Transcriptomic variation in non-radiating and radiating cichlids. (**A, B)** The absolute number of expressed isoforms identified in the oral and pharyngeal jaw mRNA of all samples. The absolute number of genes expressed is shown in Figure S4. Asterisks denote Mann-Whitney U test significance: *p* < 0.001***, *p* < 0.01**, *p* < 0.05*. NS: no significance (**C)** Percent of total genes and isoforms expressed in non-radiating species and radiations or unique to one or more of the radiations for both jaws. LV: Lake Victoria, LM: Lake Malawi, LT: Lake Tanganyika, NR: non-radiating.

### Trophic adaptation from old and new splice variants

Since the global patterns of AS revealed that it was evolving rapidly in the jaws of cichlids during adaptive radiation and that much of the splice variation was ancestral, we next asked if ancestral splice variants contributed to trophic divergence of cichlid jaws. We conducted differential PSI analysis between the jaws of species adapted to herbivorous and carnivorous diets (see Figure 1 for trophic specialisation of species) within each lake to identify differentially spliced genes (DSGs) that may play a role in shaping their eco-morphological differences. The largest number of genes were differentially spliced in Lake Victoria (OJ 303, PJ 215) followed by Lake Malawi (OJ 227, PJ 214) and Lake Tanganyika (OJ 203, PJ 126) (Figure 3A for OJ, Figure S6 for PJ, File S1). This indicates that younger cichlid radiations leveraged splice variation relatively more than the older radiation in Lake Tanganyika for trophic adaptation. Differentially spliced genes were significantly enriched (Fisher’s exact test *q* < 0.05) for GO terms specifically associated with craniofacial and jaw morphogenesis that are known from literature such as ‘regulation of osteoblasts’, ‘bone mineralisation involved in bone maturation’, ‘roof plate formation’, and ‘skeletal muscle development’ (subset shown in Figure 3C; full data in Figure S7). This was especially true for DSGs in the younger radiations. The Wnt signalling pathway that has been associated with the evolution of new craniofacial phenotypes at the larval stage in Malawi cichlids ((47) was enriched in divergent jaws from Lake Victoria. It is likely that AS of Wnt pathway genes *gpc3* and *lzts2a* (File S1) that control skeletal development and dorsoventral patterning contributed to adaptive jaw shape differences Lake Victoria cichlids ((48, 49). Two genes, *rnfox1* and *rbm5*, involved in ‘mRNA splicing via the spliceosome’ were differentially spliced in Lake Victoria oral jaws (File S1). This suggests that AS of spliceosome components, which is the molecular machinery that controls splicing, contributed to the evolution of ecologicaly flexibility in the fastest and most species-rich vertebrate adaptive radiation. In Lake Malawi OJ, differential splicing of genes involved in bone maturation (*col1a1*) and cranial morphogenesis (*sp7*) were notable. Overall, twice as many DSGs were shared between Lakes Victoria and Malawi, than either of these lakes shared with Lake Tanganyika (Figure 4A, S6), which is most consistent with these splice variants evolving and/or gaining function after the ancestor of these Lake Tanganyika haplochromines had split from the common ancestor of Lakes Victoria and Malawi, in line with the ‘Out of Tanganyika’ theory of haplochromine cichlid evolution ((27).

**Figure 4.**
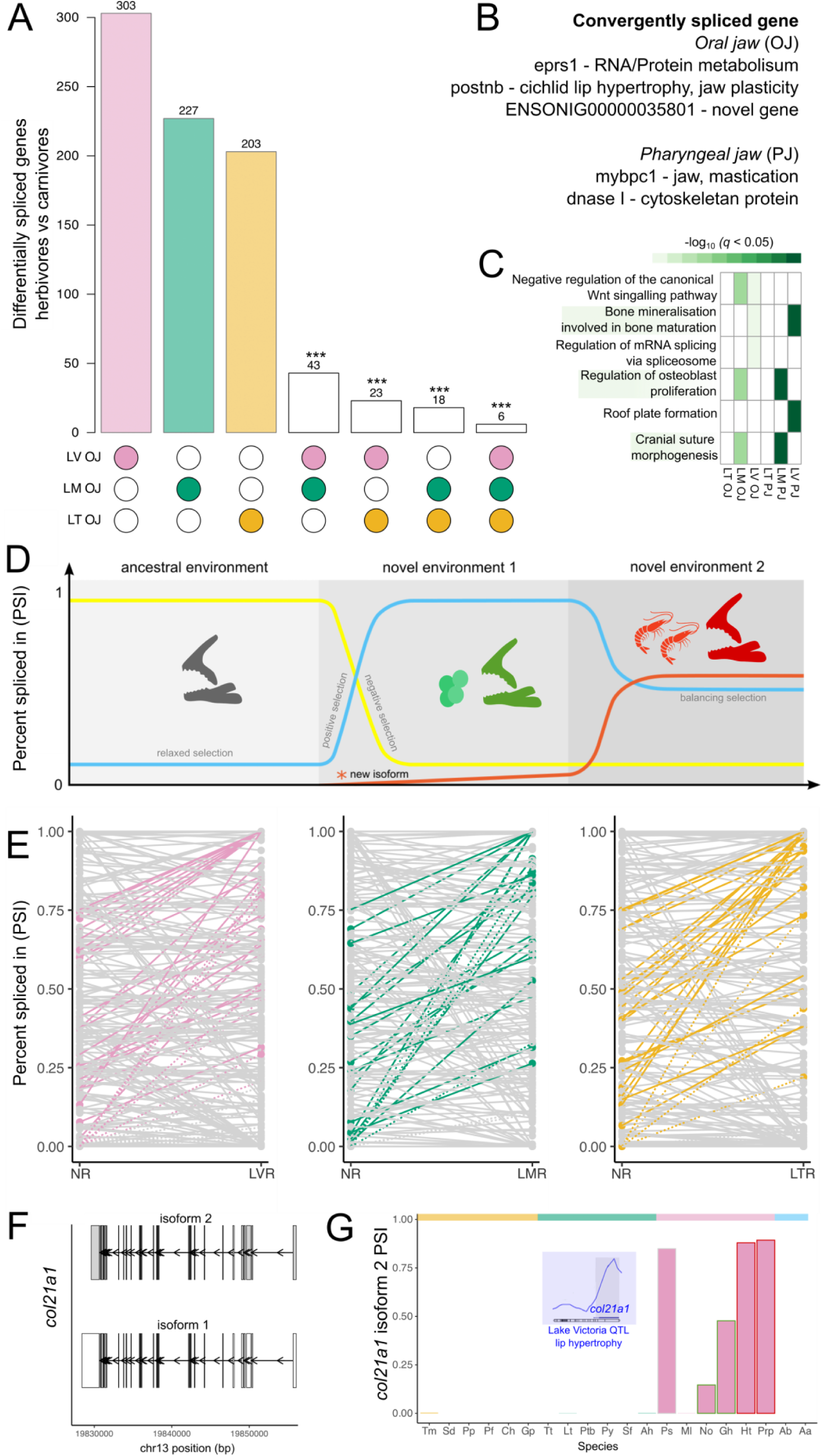
Evolutionary dynamics of differentially spliced genes between herbivores and carnivores species. (**A)** Number of significantly differentially spliced genes between oral jaws (OJ) of carnivores and herbivores within each lake radiation (*q* < 0.05) with overlaps shown (hypergeometric test *p* < 0.001). The number of differentially spliced genes between the pharyngeal jaws (PJ) of carnivores and herbivores within each radiation is shown in Figure S6A. (**B)** Genes that were convergently spliced in at least two radiations in the same direction (higher in herbivores or higher in carnivores). No gene were convergently spliced in the same directly in all three radiations. (**C)** Gene Ontology enrichment of differentially spliced genes between herbivores and carnivores. Only a subset of terms are shown. Full dataset is presented in Figure S7. (**D)** Hypothetical expectations for isoform frequency (measured as percent spliced in - PSI) during ecological adaptation. Figure adapted from Singh and Ahi (2022). (**E)** Testing the hypothesis depicted in (D) in our transcriptome data from Lake Victoria radiation (LV), Lake Malawi radiation (LM) and Lake Tanganyika radiation (LT) compared to non-radiating species (NR). Mean PSI was calculated all individuals of all species in each group (NR, LV, LM, LT). All lines represent all isoforms of genes that were signficantly differentially spliced between herbivores and carnivores within each radiation. Solid grey lines represent all isoforms that were expressd at lower or similar levels in the non-radiating species and species from each radiation (PSI differences <= 0.2). Dotted grey lines represent all isoforms that were not expressd in non-radiating species but were expressed species from each radiation (PSI differences <= 0.2). Coloured solid lines indicate isoforms that were expressed at low levels in non-radiating species but were higher expressed in radiating species (PSI differences > 0.2). Coloured dotted lines indicate isoforms that were not expressed in non-radiating species but were expressed in radiating species (PSI differences > 0.2). (**F)** The two isoform models for *col21a1* in the gene annotation. (**G)** Species mean expression of *col21a1* isoform 2 across all species in our dataset. Inset highlights a known QTL lip shape from Lake Victoria containing *col21a1*.

Of the genes that were significantly differentially spliced between herbivores and carnivores in each lake (Figure 3A, S6A), a dynamic pattern of isoform evolution emerged: many of these were low expressed isoforms in non-radiating species that increased in expression in radiating species (Figure 3E for OJ, S6B for PJ). An increase in frequency would be the expectation if these low expressed ancestral alternative isoforms persisted in populations as standing variation due to relaxed selection pressures (38, 39) and gained adaptive function when these lineages entered lakes and discovered newly available food sources (Figure 3D) as hypothesised by Singh & Ahi ((5). Many highly expressed isoforms in non-radiating species also decreased in frequency or were lost in the lacustrine radiations (presumably as the alternative isoform increased in frequency). We further identified 34 potentially novel isoforms that were absent in non-radiating species and only expressed a subset of the three adaptive radiations (Figure 3E dotted lines, File S1). Sixteen of these 34 novel isoforms were found in LV, followed by 12 in Lake Malawi and six in Lake Tanganyika. Novel isoforms of note were of genes *col21a1* and *lzts2a*. *Col21a1* had two isoforms (Figure 3F) of which the main isoform is expressed in the OJ of Lake Victoria herbivores as well as herbivores from Lakes Tanganyika and Malawi (with one exception, File S1). The alternative isoform of *col21a1* is only expressed in Lake Victoria species, with higher expression in carnivores and omnivores (Figure 3G). *Col21a1* is associated with human cleft lip syndrome and maps to a QTL for hypertrophic lips in Lake Victoria cichlids(50). It is conceivable that this isoform may have arisen *de novo* in Lake Victoria cichlids or arrived through hybridisation, which has been shown to contribute genetic variation for rapid adaptive radiation in Lake Victoria(44). In the future, integrating transcriptomic analysis with population genetics and functional genomics (F_ST_ scans, GWAS, spliceQTL) approaches would help to narrow down the genetic variants associated with splicing and the evolutionary processes that shape them. Our results also highlight the conservation of craniofacial development pathways across vertebrates, and the promise of cichlid fishes as a natural model for studying human craniofacial disease(51).

### Differential expression of major craniofacial genes contributed to jaw divergence

In addition to differences in splicing, we investigated GE differences between herbivores and carnivores in each lake radiation to be able to compare the role of GE with AS in trophic adaptation. The largest number of differentially expressed genes (DEGs) were in Lake Tanganyika (OJ 5506, PJ 4094), followed by Lake Malawi (OJ 1970, PJ 1499) and Lake Victoria (OJ 1792, PJ 1166) (Figure 5A for OJ, Figure S8 for PJ, File S1). Several interesting candidate genes for jaw/craniofacial morphogenesis known from literature stood out as the most extreme outliers in the oldest Lake Taganyika where GO terms such as ‘neural crest cell development’, ‘skeletal system development, and ‘ossification’ were significantly enriched (*q* < 0.05). One such gene for Lake Tanganyika OJ was Frizzled 2 (*fzd2*), a Wnt pathway gene that is highly expressed in the developing face of vertebrates, and mutations of this gene cause cleft palate and wider nose and upper jaw(52). A gene controlling prey crushing teeth morphology in cichlids, *odam (*(53), and a gene linked to craniofacial disease, *gja1,* were also significant for Lake Tanganyika OJ(54). Two interesting outliers in Lake Victoria OJ were *kaznb*, a craniofacial development gene that has been found to be upregulated in the jaws of herbivorous cichlids at larval and adult life stages (55) and *egr1*, that forms part of the regulatory cascade modulating cranial bone morphogenesis through the BMP pathway(56). Interestingly, one of the most extreme outliers for Lake Taganyika OJ was *hnRNPk*, an RNA binding protein that regulates pre-mRNA splicing(57). Overall, the GO terms for DEGs in the youngers lakes were less specific to jaw development than the DSGs. This suggests that shifts AS may be a more flexible route to rapid diversification in explosive adaptive radiations than changes in GE.

**Figure 5.**
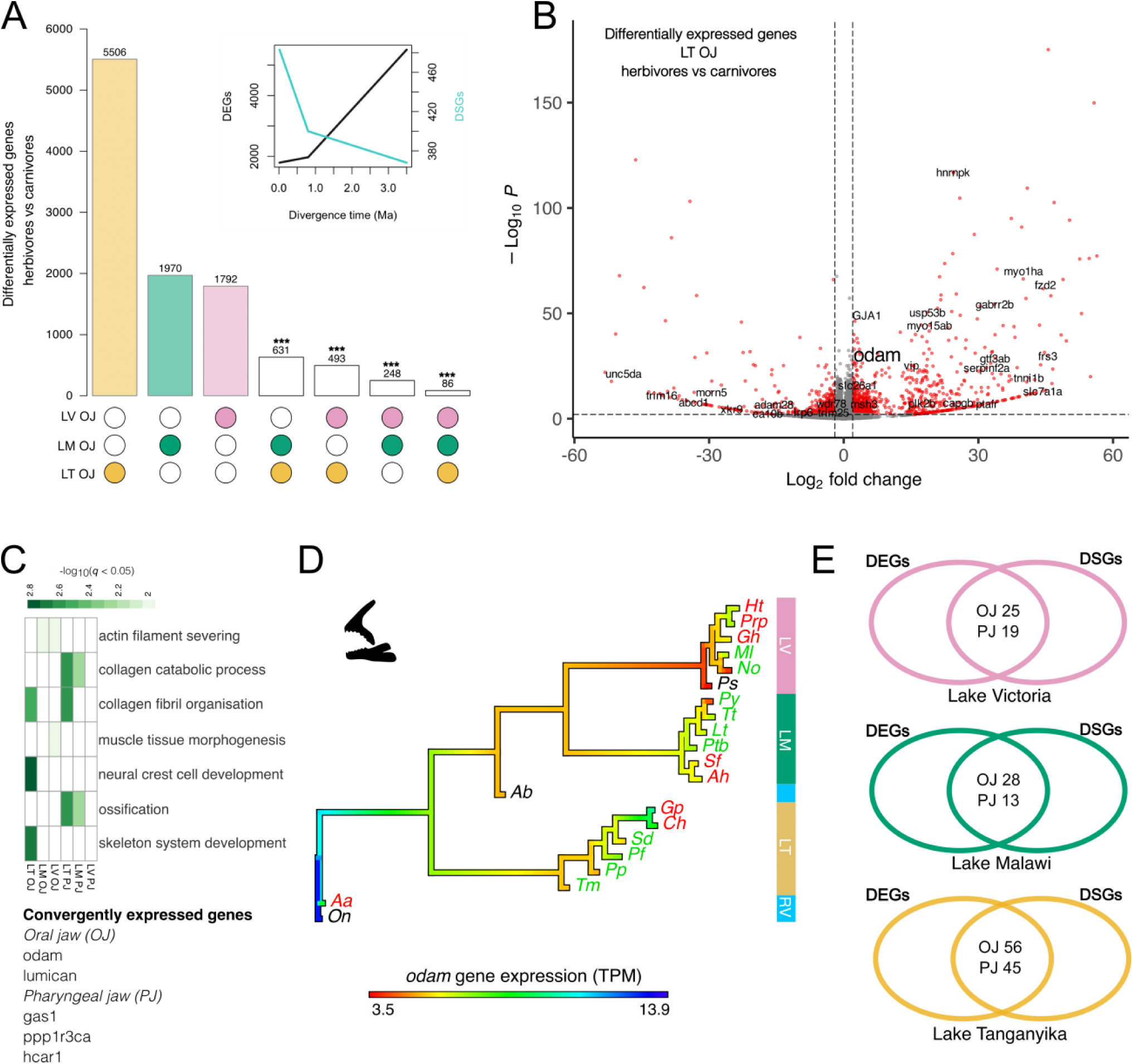
GE divergence in oral jaws of herbivore and carnivore species. **(A)** Number of differentially expressed genes (DEGs, *q* < 0.05) between oral jaws (OJ) of herbivores and carnivores in each lake, with overlaps annotated (hypergeometric test *p* < 0.001). Results for pharyngeal jaws can be found in Figure S8. Inset shows the relationships between number of DEGs and differentially spliced genes (DSGs) in each cichlid radiationnplotted as function of divergence time. (**B)** Volcano plot illustrating DEGs in Lake Tanganyika (LT) oral jaws. Significant DEGs are highlighted in red. Top gene outliers are annotated. (**C)** Gene Ontology enrichment of DEGs between herbivores and carnivores. Only a subset of terms are shown that are known in literature to be associated with jaw/craniofacial divergence. Full list is presented in Figure S9. Some candidate genes for jaw morphogensis/craniofacial development that were convergently expressed in all three radiations in the same direction (higher in herbivores or higher in carnivores). See File S1 for full list. (**D)** Ancestral state reconstruction of GE of the convergently expressed *odam* gene. (**E)** Overlap of DEGs and DSGs between herbivores and carnivores within each lake radiation.

### Contrasting functions for alternative splicing and gene expression

We found dissimilar patterns in the extent of differential GE and AS underlying divergent trophic adaptation across cichlid radiations. Even though the overall number of DEGs was much higher than DSGs across all three radiations (Figures 4A, 5A, S6, S8), the relative number of DEGs across radiations was the highest in the oldest radiation and lowest in the youngest radiation, while the DSGs showed the opposite pattern (Figure 5A inset). Interestingly, more DEGs in each of the younger radiations overlapped with the older radiation, in contrast more DSGs overlapped among the younger radiations (Figures 4A, 5A, S6A, S8). This pattern indicates that a complex temporal interplay of GE and AS underlies adaptive radiation, with ancestral splice variation enabling rapid ecological diversification at early stages of speciation, while GE differences evolve much slower, mostly likely fine-tuning the adaptive morphology long after speciation. This would be consistent with the GO enrichment results for differentially spliced and expressed genes that we found in younger versus the older radiation.

It has been proposed that AS and GE perform contrasting biological functions(11, 55, 58–60). However, these studies have lacked a phylogenetic perspective. In our dataset, the overlap of genes that were both differentially spliced and differentially expressed between herbivores and carnivores followed a phylogenetic and evolutionary pattern. The highest overlap was in Lake Tanganyika (OJ 56, PJ 45), followed by Lake Malawi (OJ 28, PJ 13) and Lake Victoria (OJ 25, PJ 19) (Figure 5). Our results reinforce the idea that putatively adaptive AS and GE changes act mostly at different genes – but we show that the extent of gene overlap depends on the evolutionary age of lineages, with fewer genes being regulated by both procesess in more recently diverged populations.

### Convergent gene expression and splicing underlying convergent trophic evolution

African cichlid radiations are an ideal sytem to study the repeatability of evolution as they have evolved convergent trophic adaptations, particularly in jaw shape, to exploit similar food sources in similar ways in different lakes(19). See results of geometric morphometric analysis in the Supplementary material that illustrates that herbivores and carnivores cluster together in a jaw shape PCA regardless of which lake they are from; and that jaw shape is diverging along parallel axes in the three radiations. We asked if differentially expressed and spliced genes contributing to herbivory vs carnivory showed parallel divergence across all three radiations. We find convergent DEGs in both oral (23 upregulated, 12 downregulated in herbivores) and pharyngeal jaws (14 upregulated, 6 downregulated in herbivores) that fit this pattern (Figures 5, S8). Several convergently used OJ genes have been associated with the development of teeth (*odam*), bone (*lum*) and muscle (*myrip*, *myo15aa*) (Figure 5B,C, File S1). *Odam* expression has been associated with larger vs smaller teeth in cichlid fishes and our ancestral state reconstruction suggests that this gene increased in expression repeatedly in the oreal jaws of carnivores in each lake radiation (Figure 5D). Convergent genes for PJ were *hcar1*, *ppp1r3c*, myl13, and *gas1*. *Gas1* interacts with *shh* is early face development (61). We validated the convergent expression of *odam, gas1, hcar1,* and *ppp1r3ca* using qPCR (Figure S10). No DSGs were convergently spliced in the same direction in all three radiations. We only found evidence of convergent splicing in the same direction in two of the three radiations in the following noteworthy genes: OJ: *eprs1*, *postnb* and PJ: *mybpc1*, *Dnase1* (Figure 4A, B). These results suggest that jaw adaptation in cichlids is a polygenic trait with different genes functioning redundantly across these three radiations. The redundancy is higher in the regulation of genes through AS than GE, most likely due to the species-specific nature of AS.

## Discussion

Bursts of diversification that characterise adaptive radiations have challenged the expectations of gradual evolution (62, 63). This divergence is often faster than the rate at which protein-coding mutations accumulate, making gene regulation an important mediator of rapid and diverse phenotypic change (3, 4, 8). However, we still understand little about how different gene regulatory mechanisms shape rapid adaptive radiations (5). Through the analysis of 200 transcriptomes representing three independent but phylogenetically nested cichlid adaptive radiations that evolved in the last <16,000 to 3.8 million years we show that the rich cache of ancestral isoform variation in the ancestor of East African haplochromine cichlids evolved faster than gene expression to rapidly to give rise to novel trophic adaptations. This pattern was most pronoucned in the youngest and most species rich adaptive radiations in Lakes Victoria and Malawi that had releatively higher differences in splicing than Lake Tanganyika of genes are associated with jaw morphogenesis. We further discovered that novel isoforms that most likely evolved in the course of adaptive radiation also contributed to adaptation. The rapid turnover of AS that we observed here is consistent with relaxed selection pressures on splice-sites, as alternative isoforms can escape the effects of purifying selection as long as the major isoform is expressed (38). Theory suggests that relaxed selection can increase the rate of evolution of complex adaptations in heterogenous environments, allowing populations to rapidly cross adaptive valleys (64). We propose that relaxed selection maintaining standing splice variation, followed by periods of positive selection on alternative isoforms of craniofacial genes contributed to the rapid evolution of trophic adaptations in cichlids as they entered lakes and were exposed to new and ecologically more stable environments. An alternative explanation for our observation can be the complex influence of diverse forms of selection (perhaps in combination with neutral processes), such as balancing selection, frequency-dependent selection, and diversifying selection, that promote isoform diversity. Taken together, our findings challenge the long-held notion that most alternative isoforms are evolutionary noise (65), because this noise can fuel innovation when ecological opportunity arises, providing one possible mechanistic explanation for how these extraordinary cichlid radiations were able to exploit ecological opportunity as such extremely short timescales.

Our results on the evolutionary dynamics of GE are partially consistent with previous work on transcriptome evolution at deeper timescales. In contrast to AS, GE is largely conserved at the tissue-level during vertebrate evolution (40, 41, 66, 67). Remarkably, we found lineage-specific clustering of GE in Tropheini jaws. Tropheini are named after their diverse trophic adaptations that allowed them to enter the littoral zones of Lake Tanganyika at a much later stage and successfully diversify alongside other established cichlid radiations(26, 68, 69). They were able to do this by rapidly remodelling their jaw, dentition, and craniofacial morphology to forage foods in new and different ways (68). Our findings may reflect the particular importance of gene regulation for evolutionary transitions in Tropheini jaws during speciation. (74)

We find that signatures of splicing became increasingly more species-specific, and GE became more lineage-specific over evolutionary time. However, specific-specific signatures of splicing do not emerge until ∼3.8 million years of evolution, a pattern not observed in previous large-scale studies on older lineages (40, 66). We also found that both rapidly evolving AS and GE contributed to jaw adaptations in herbivorous and carnivorous cichlids, but their contribution was inversely predicted by evolutionary age. Relatively more differences in AS were found to underlie trophic differences in jaws of cichlids from the younger radiations, while GE differences dominated in the older radiation. This was also reflected in the biological functions of these genes: (1) more jaw morphogenesis related gene ontology terms were enriched in differentially spliced genes in Lakes Victoria and Malawi than Tanganyika; (2) in Tanganyika more jaw morphogensis related gene ontology terms were enriched in differentially expressed genes. This pattern indicates that a complex temporal interplay of GE and AS underlies adaptive radiation, with ancestral splice variation enabling rapid ecological diversification at early stages of speciation, while GE differences evolve much slower, mostly likely fine-tuning the adaptive morphology long after speciation. Consistent with previous studies, we found many genes implicated in jaw divergence suggesting that jaw remodelling is a complex phenotype governed by myriad genes (15, 19, 26, 70).

The Wnt pathway emerged as a key player in the evolution of new jaw phenotypes at early stages of development, with genes regulated through splicing (*gpc3*, *lzts2a*) and differential expression (*fzd2*). Overall, more convergent differences in GE than AS were found to underlie convergen trophic jaw adaptations across all three cichlid radiations. The lower phylogenetic convergence of AS may underscore its role in shaping functional genetic redundancy - where multiple genetic pathways can produce similar phenotypic outcomes (71). In our case, it is not isoform diversity within a single gene that underpins redundancy as classically discussed (72) but rather the use of different genes regulated by lineage-specific splicing patterns, highlighting a novel form of genetic redundancy mediated by AS. Such redundancy may allow different lineages to evolve comparable traits via distinct molecular routes, increasing evolutionary flexibility and speed.

It has been hypothesised that ancestral variation plays an important role in the rapid adaptive capacity of African cichlids (31, 73). But, little empirical evidence exists explaining how this ancestral variation mechanistically plays out. We show that ancestral splice variants that were differentially spliced between jaws of herbivores and carnivores are lowly expressed in non-radiating species but increased in frequency in radiating cichlids, most likely enabling trophic divergence. Firstly, this suggests that old genetic variation can fuel new adaptations (31); and that the ancestors of the East African cichlid adaptive radiations, which were most likely a widely dispersing riverine cichlids, acted as reservoirs of genetic variation for diversification on a continental scale (74). And secondly, even if most ancestral alternative isoforms are non-functional (65), lowly expressed, and evolving stochastically (38), they represent a cache of standing variation for adaptive evolution when ecological opportunity arises (5, 75). Standing variation in AS has been shown to be important for the rapid domestication of sunflowers (75), which represents only one ecological transition, while cichlid adaptive radiations involve multiple, often convergent, diverse ecological transitions and speciation events. In addition to ancestral variation, we found evidence of potentially novel isoforms contributing to adaptive jaw differences. This emphasises how fast new protein-coding variation can be generated for evolution through AS.

Our present study is one of the most comprehensive cross-lake analysis of transcriptomic variation in cichlid fishes to date. However, it stills represents a small subset of the hundreds of cichlid species that are found in the African Great Lakes. In the future, expanding the analyses to include more species would add more confidence to our findings on convergent and divergent gene expression and splicing patterns underlying trophic evolution in these radiations. Furthermore, including more cichlid species that entered lakes and had access to ecological opoortunity but did not form radiations (76), or riverine cichlids that are hypothesised to have contributed genetically to the lake radiations through hybridisation (25, 44, 45), would shed light on how different evolutionary processes promote (or constrain) rapid diversification by re-wiring transcriptional variation. Targeting a single, albeit ecologically and morphologically important, developmental stage is another constraint of our approach. In the future, studying gene regulation across multiple developmental stages, in combination with how jaw shape changes along ontogeny, would really enhance our understanding of gene regulation and the origins of diversity.

Taken together, our findings suggest that AS may be one piece of the puzzle explaining how ecological and species richness can evolve when sufficient time is not available for new protein-coding mutations to arise, fix, and be replenished. Future studies on gene regulation in evolutionary biology would benefit from moving away from a gene expession-centric view to a more holistic model that incoporates other levels of gene regulation, such as splicing.

## Materials and Methods

### Study design and sampling

In this study we focused on six cichlid species from Lake Tanganyika, six cichlid species from Lake Malawi, six cichlid species from Lake Victoria as well as two cichlid species from surrounding rivers, all of which are part of the ‘modern haplochromine’ adaptive radiations (Figure 1). The 18 species from the three lakes represent eco-morphological replicates of herbivore and carnivore trophic adaptations. We focused on the functionally decoupled oral (OJ) and pharyngeal jaw (PJ) tissues, that are involved in food capture and processing.

Sample were collected by either through aquarium trade or fieldwork within the framework of collaborations with Zambian and Tanzanian fisheries (TAFIRI). All fish were reared in the fish facility at University of Graz under standardised conditions regarding tank environment, diet, and water parameters to minimise plastic effects. Fertilised eggs were taken from the mouth of mouthbrooding females 24 hours after fertilisation, by applying slight pressure onto their cheeks. The eggs were reared in a separate tank until the yolk sac was completely absorbed, which is also known as the developmental stage 26 – end of larval stage and beginning of juvenile stage, when bony elements have formed but the organism is still growing actively (77–79). Late larval and juvenile developmental stages are ecomorphologically informative stages that have been used to study jaw adaptation and divergence cichlids (11, 80, 81). The external morphology and internal bone and cartilage development at stage 26 is illustrated in Fig. S11. Stage 26 larvae were euthanised and stored in RNAlater in a fridge for seven days and then moved to a -80°C freezer. (11, 15, 72)

### RNA extraction

Each larva was dissected under a stereo microscope to dissect the oral jaws (OJ: bone, teeth) and the pharyngeal jaws (PJ: bones and teeth) separately (Figure S12). The bones were cleaned as much as possible, but some surrounding tissue (i.e. skin, cartilage, muscle) remained, especially for the OJ, as small size of the larvae (<1cm) makes the dissection challenging. The dissected tissue was submerged in a lysis buffer and 1-thioglycerol according to the protocol of the Promega RNA Reliaprep kit. The sample in the lysis buffer was crushed with a ceramic bullet in a homogenizer (FastPrep-24, MP Biomedicals, Santa Ana, CA, USA) before further processing. Two individuals were pooled per sample to increase the amount of RNA for Lake Victoria species. For Lake Malawi and Tanganyika, one individual yielded sufficient RNA. RNA quality was tested at a TapeStation 2200 with RNA Screen Tapes. It was tried to reach a RINe number of >= 7 for each sample. We extracted RNA for 5 biological replicates per jaw per species, which resulted in 200 mRNA samples in total.

### Library preparation and sequencing

Library preparation was done with the TruSeq Stranded mRNA Sample Prep Kit using 24 indexing adapters and following the standard protocol. For the input RNA we tried to reach 1000ng for each sample. The quality of the cDNA was tested on a TapeStation 2200 with D1000 Screen Tapes. Sequencing was executed at the Vienna BioCenter Core Facilities on a HiSeq 2500 with paired end 125 cycles per read (2 x 125 cycles). Demultiplexing was conducted by the same facility.

### Transcriptome assembly

Each sample has approximately ten million paired-end reads. Raw sequence reads are available at NCBI sequence read archive (Reviewer link: https://dataview.ncbi.nlm.nih.gov/object/PRJNA640176?reviewer=p2dhiiern60seqodpuqbqb1o62). After a quality check with Fastqc (v0.11.8)(*82*) and a trimming step with Trimmomatic (v0.3.9)(83), only reads with a phred > 28 and a minimum length of 70 bp where retained. To enable the assembly of novel isoforms we developed a bioinformatics pipeline with the following steps: Reads were assembled using STAR (v2.7.3.a)(84) in reference guided mode using the *Oreochromis niloticus* reference genome(*85*). This genome was selected because it is the best assembled and annotated cichlid reference genome and is phylogenetically equidistant to the species sampled in this study. Mapping statistics were generated with samtools idxstats (v1.9)(33, 86). StringTie (v2.0.6) was run first both with a reference genome to assemble the RNA-Seq alignments into potential transcripts and in genome-guided mode. This was done separately for the single files (per individual). The single gtf files for each biological replicate were then incrementally merged according to species and tissue and finally into one super annotation. This repeated merging steps were conducted to reduce the probability of false positives in isoform assembly and accurately identify novel species-specific or lower expressed isoforms. To estimate the accuracy of the produced annotation files we compared them with gffcompare (v0.11.2)(*87*) to our reference *O. niloticus* annotation. We filtered for monoexonic transcripts not contained in the reference and a class code assigned by gffcompare (v0.11.2), indicating ‘possible polymerase run-on’ fragments. The maximum intron length was reduced to 200,000 bp, which is the maximum intron length found in the *O.niloticus* reference. Based on the super annotation, the expression estimates were generated with StringTie allowing no multimapping. From these expression estimates, count matrices were produced using a perl script from the griffith lab (https://rnabio.org) to extract raw count data from StringTie results. The code used for our RNAseq pipeline in on github(88). SUPPA2 (v2.3)(37) was used to calculate percent spliced in (PSI) index for each isoform for each gene.

### Differential gene expression analysis

We used DESeq2(34)in R(89) to detect DEGs running comparisons (1) between herbivores and carnivores within each lake radiation for each tissue separately and (2) between oral and pharyngeal jaws within each radiation. DESeq2 uses raw read counts and estimates variance-mean dependence based on a model that utilises the negative binomial distribution. The cutoff for DEGs was chosen at a false discovery rate of (*q* < 0.05).

### Alternative splicing analysis

Alternative splicing analysis was conducted with SUPPA2 (v2.3)(37) to evaluate differential transcript usage (DTU) between between herbivores and carnivores within each lake radiation for each tissue separately. Visualisation of PSI analysis was conducted in R(89).

### Modeling gene and isoform expression

We used the methods proposed by Chen et al (2019)(42) to model gene and isoform expression along the phylogeny using Ornstein-Uhlenbeck (OU) and Brownian Motion (BM) processes. The OU process is a modification of a random walk, describing the change in expression (dX_t_) across time (dt) by dX_t_ = σdB_t_ + α (θ – X_t_) dt, where dB_t_ denotes a Brownian motion process. The model elegantly quantifies the contribution of both drift and selective pressure for any given gene: (1) Drift is modeled by Brownian motion with a rate σ while (2) the strength of selective pressure driving expression back to an optimal expression level θ is parameterized by α. The OU process accounts for phylogenetic relationships, thus allowing us to fit individual evolutionary expression trajectories.

### Phylogenetic analysis and ancestral state reconstruction of gene expression

We constructed expression trees using the neighbour-joining approach based on pairwise distance matrices between samples using data from all genes and applying functions from the ’ape’ package (v5.8.1)(90) in R(89). Distances were computed as 1 − ρ, where ρ is the Spearman’s correlation coefficient, chosen due to its robustness against outliers and any normalisation issues(41). To visually evaluate the differences in expression evolution, we constructed a tanglegram of the expression phylogenies of the two jaw tissues using the ‘dendextend’ package (v1.19.0)(91) in R. Ancestral state reconstruction was conducted using the ‘phytools’ package (v2.4.4)(92) in R.

### Phylogenetic reconstruction (SNPs)

SNP calling on mapped mRNAseq reads was conducted using bcftools(93) across all samples. Filtered high quality SNPs were used to contruct a concatenated alingment with the most common allele at each position for each individual. This alignment was used as input for RAxML (v8)(94) to contstruct a SNP phylogeny.

### GO term enrichment

GO terms for *Oreochromis niloticus* were acquired in R via the biomaRt package (v2.46.1)(95) from ensembl(96). Gene set enrichment analysis was conducted using topGO (v2.36.0)(*97*) using the method weight to account for GO topology and Fisher’s exact test to correct for multiple testing for the enrichment analysis. We used our super annotation to build the gene universe.

(98)(2)(98, 99)(98, 99)

## Data Availability

All data has been deposited to the NCBI SRA BioProject (PRJNA640176). Reviewer’s link: https://dataview.ncbi.nlm.nih.gov/object/PRJNA640176?reviewer=p2dhiiern60seqodpuqbqb1o62 All bioinformatics code can be found on github: https://github.com/poojasingh09/2024_singh_transcriptomeanalysis_pipeline

## Notes

### Competing Interest Statement

The authors have declared no competing interest.

### Summary of Updates

Limitations section added Geometric morphometric analysis added

### Funding

- FWF Austrian Science Fund

