## Supplementary figures and tables for "Ancestral splice variation is a key substrate for rapid diversification in African cichlids"

Pooja Singh *et al*

Supplementary Information

**Table S1 Species information**

| Abbreviation | species | Trophic niche |
|---|---|---|
| | **Lake Malawi** | |
| Ah | *Aulonocara hansbaenschi* "Red Flush" | carnivore (invertebrates) |
| Lt | *Labeotropheus trewavasae* "Red Top Thumbi West" | herbivore (algae) |
| Ptb | *Petrotilapia* sp. "Thick bars" | herbivore (algae) |
| Py | *Petrotilapia* sp. "Yellow chin Chewere" | herbivore (algae) |
| Sf | *Sciaenochromis fryeri* | carnivore (small fish, invertebrates) |
| Tt | *Tropheops tropheops* "Makokola" | (algae, plankton) |
| | **Lake Tanganyika** | |
| Ch | *Ctenochromis horei* | carnivore (small fish, invertebrates) |
| Gp | *Gnathochromis pfefferi* | carnivore (invertebrates) |
| Pf | *Petrochromis famula* "Crocodile Island" | herbivore (algae, phytoplankton) |
| Pp | *Petrochromis polyodon* "East Zambia" | herbivore (algae) |
| Sd | *Simochromis diagramma* | Herbivore (algae, phytoplankton) |
| Tm | *Tropheus moorii* "Nakaku" | Herbivore (algae, phytoplankton) |
| | **Lake Victoria** | |
| Gh | *Gaurochromis hiatus* 'Mwanza Gulf' | carnivore (invertebrate) |
| Ps | *Paralabidochromis sauvagei* (Haplochromis sp. 'rock kribensis') | omnivore (invertebrate, algae) |
| Ht | *Haplochromis thereuterion* | carnivore (invertebrate, zooplankton) |
| Ml | *Mbipia lutea* ' Makobe island' | herbivore (algae) |
| No | *Neochromis omnicaeruleus* "Makobe Island" | herbivore (algae) |
| Prp | *Prognathochromis perrieri* | carnivore (small fish) |
| | **Non-radiating riverine cichlids that also entered lakes** | |
| Aa | *Astatoreochromis alluaudi* | carnivore (invertebrates) |

| | | |
|---|---|---|
| Ab | *Astatotilapia burtoni* "chipwa" | omnivore (algae, invertebrates, fish remains, opportunistic feeder) |

**Table S2 RNA mapping statistics**

| Category | Uniquely mapped | Mapped to multiple loci | Mapped to too many loci | Unmapped: too short | Unmapped: other | Uniquely mapped (%) |
|---|---|---|---|---|---|---|
| Aa.OJ.1 | 5115846 | 319640 | 37437 | 922391 | 96069 | 78.8 |
| Aa.OJ.2 | 4130953 | 273830 | 70124 | 785922 | 221024 | 75.4 |
| Aa.OJ.3 | 5563207 | 352615 | 28881 | 1071263 | 54447 | 78.7 |
| Aa.OJ.4 | 22083577 | 1376678 | 124565 | 4003993 | 244859 | 79.3 |
| Aa.OJ.5 | 6853276 | 433792 | 38157 | 1279754 | 77271 | 78.9 |
| Aa.PJ.1 | 5261149 | 325804 | 85758 | 835943 | 287483 | 77.4 |
| Aa.PJ.2 | 3728025 | 295212 | 173541 | 750399 | 680595 | 66.2 |
| Aa.PJ.3 | 4317572 | 237349 | 25333 | 737273 | 54274 | 80.4 |
| Aa.PJ.4 | 4182693 | 256788 | 28696 | 654894 | 62223 | 80.7 |
| Aa.PJ.5 | 4410212 | 276925 | 27654 | 794798 | 60153 | 79.2 |
| Ab.OJ.1 | 4188515 | 256459 | 25177 | 685525 | 60472 | 80.3 |
| Ab.OJ.2 | 5488288 | 322164 | 27270 | 927459 | 54556 | 80.5 |
| Ab.OJ.3 | 4402449 | 253917 | 22692 | 703783 | 49417 | 81 |
| Ab.OJ.4 | 5953687 | 347155 | 28348 | 1094816 | 58370 | 79.6 |
| Ab.OJ.5 | 4979488 | 276587 | 21549 | 914409 | 39865 | 79.9 |
| Ab.PJ.1 | 2760915 | 154480 | 21365 | 421941 | 46655 | 81.1 |
| Ab.PJ.2 | 6501582 | 379333 | 64527 | 958128 | 148148 | 80.7 |
| Ab.PJ.3 | 4795157 | 259492 | 30924 | 645173 | 64343 | 82.7 |
| Ab.PJ.4 | 3510209 | 189632 | 16834 | 504030 | 32326 | 82.5 |
| Ab.PJ.5 | 3038370 | 179106 | 23621 | 486109 | 44500 | 80.6 |
| Ah.OJ.1 | 5307986 | 320283 | 25064 | 1080552 | 42680 | 78.3 |
| Ah.OJ.2 | 6405151 | 389208 | 31086 | 1335849 | 51726 | 78 |
| Ah.OJ.3 | 7154864 | 451190 | 50059 | 1401315 | 107229 | 78.1 |
| Ah.OJ.4 | 6004083 | 358058 | 31666 | 1182590 | 58022 | 78.6 |
| Ah.OJ.5 | 4680805 | 271972 | 22633 | 896581 | 41372 | 79.2 |

| | | | | | | |
|---|---|---|---|---|---|---|
| Ah.PJ.1 | 4818647 | 291360 | 23631 | 886664 | 40000 | 79.5 |
| Ah.PJ.2 | 4135397 | 261319 | 25389 | 781787 | 45679 | 78.8 |
| Ah.PJ.3 | 5348725 | 316474 | 23864 | 1047911 | 42703 | 78.9 |
| Ah.PJ.4 | 4520939 | 262270 | 23939 | 807678 | 45280 | 79.9 |
| Ah.PJ.5 | 4168385 | 241142 | 25891 | 697131 | 50276 | 80.4 |
| Ch.OJ.1 | 5002452 | 262198 | 26178 | 1056652 | 64798 | 78 |
| Ch.OJ.2 | 2497732 | 125361 | 14636 | 449960 | 28025 | 80.2 |
| Ch.OJ.3 | 3829827 | 205598 | 19696 | 839198 | 49452 | 77.5 |
| Ch.OJ.4 | 4444873 | 236075 | 21130 | 798869 | 50519 | 80.1 |
| Ch.OJ.5 | 4034325 | 206288 | 20026 | 705193 | 47113 | 80.5 |
| Ch.PJ.1 | 3479115 | 170941 | 17532 | 568334 | 38031 | 81.4 |
| Ch.PJ.2 | 5992763 | 304085 | 27964 | 1619671 | 58417 | 74.9 |
| Ch.PJ.3 | 5463718 | 268809 | 22711 | 848127 | 51220 | 82.1 |
| Ch.PJ.4 | 5498193 | 269773 | 24618 | 934110 | 63134 | 81 |
| Ch.PJ.5 | 4588238 | 227873 | 18698 | 791605 | 45964 | 80.9 |
| Gh.OJ.1 | 5852007 | 335325 | 32142 | 1040482 | 53377 | 80 |
| Gh.OJ.2 | 3667558 | 205744 | 22582 | 781410 | 36761 | 77.8 |
| Gh.OJ.3 | 7658391 | 426771 | 35351 | 1352469 | 60089 | 80.3 |
| Gh.OJ.4 | 5957258 | 349157 | 27034 | 1130333 | 47316 | 79.3 |
| Gh.OJ.5 | 7008441 | 408006 | 29622 | 1368109 | 47848 | 79.1 |
| Gh.PJ.1 | 3867659 | 203217 | 22476 | 760651 | 35686 | 79.1 |
| Gh.PJ.2 | 4719313 | 253932 | 30370 | 933675 | 47881 | 78.9 |
| Gh.PJ.3 | 4460176 | 231912 | 25418 | 876301 | 38296 | 79.2 |
| Gh.PJ.4 | 5352295 | 292364 | 23408 | 1074392 | 42080 | 78.9 |
| Gh.PJ.5 | 4717261 | 257892 | 22559 | 983354 | 40948 | 78.3 |
| Gp.OJ.1 | 6022120 | 306589 | 14912 | 1193898 | 42450 | 79.4 |
| Gp.OJ.2 | 4757192 | 228283 | 14902 | 1189862 | 36755 | 76.4 |
| Gp.OJ.3 | 4928780 | 237200 | 15344 | 1261352 | 38242 | 76.1 |
| Gp.OJ.4 | 3924503 | 189202 | 14650 | 1011090 | 39366 | 75.8 |
| Gp.OJ.5 | 5472749 | 289954 | 84381 | 1013191 | 283665 | 76.6 |
| Gp.PJ.1 | 5001575 | 269485 | 18017 | 1051661 | 52423 | 78.2 |
| Gp.PJ.2 | 4427633 | 218902 | 19156 | 977264 | 60473 | 77.6 |
| Gp.PJ.3 | 3686019 | 178769 | 23781 | 885692 | 61399 | 76.2 |
| Gp.PJ.4 | 4123711 | 217958 | 54869 | 970168 | 195814 | 74.1 |
| Gp.PJ.5 | 2549939 | 129874 | 29799 | 443173 | 76231 | 79 |
| Ht.OJ.1 | 6911297 | 424070 | 33186 | 1458875 | 61305 | 77.8 |

| | | | | | | |
|---|---|---|---|---|---|---|
| Ht.OJ.2 | 3897780 | 229901 | 42169 | 771468 | 66083 | 77.8 |
| Ht.OJ.3 | 6590962 | 361908 | 23565 | 1227627 | 38750 | 80 |
| Ht.OJ.4 | 4482925 | 228603 | 26133 | 864845 | 41747 | 79.4 |
| Ht.OJ.5 | 7182555 | 418861 | 34804 | 1368159 | 58894 | 79.2 |
| Ht.PJ.1 | 3208610 | 178816 | 13082 | 701154 | 22684 | 77.8 |
| Ht.PJ.2 | 2307733 | 135142 | 30230 | 468787 | 57286 | 76.9 |
| Ht.PJ.3 | 5839520 | 303298 | 23626 | 1179712 | 44350 | 79 |
| Ht.PJ.4 | 2976068 | 152394 | 17593 | 552974 | 30223 | 79.8 |
| Ht.PJ.5 | 3275466 | 174130 | 18598 | 600324 | 30733 | 79.9 |
| Lt.OJ.1 | 5592178 | 298660 | 25073 | 1083286 | 50746 | 79.3 |
| Lt.OJ.2 | 5466662 | 303939 | 22271 | 993950 | 38203 | 80.1 |
| Lt.OJ.3 | 7169613 | 400601 | 30072 | 1440579 | 49111 | 78.9 |
| Lt.OJ.4 | 6231704 | 353789 | 27106 | 1185504 | 47860 | 79.4 |
| Lt.OJ.5 | 6542186 | 363235 | 28388 | 1177952 | 55510 | 80.1 |
| Lt.PJ.1 | 3848024 | 191386 | 16838 | 768084 | 39401 | 79.1 |
| Lt.PJ.2 | 4994867 | 269158 | 16761 | 934318 | 29978 | 80 |
| Lt.PJ.3 | 6274784 | 344150 | 23467 | 1137165 | 47741 | 80.2 |
| Lt.PJ.4 | 4111017 | 230908 | 15048 | 799248 | 25916 | 79.3 |
| Lt.PJ.5 | 5088894 | 260944 | 19563 | 851570 | 44490 | 81.2 |
| Ml.OJ.1 | 4779528 | 271544 | 21486 | 982068 | 37772 | 78.5 |
| Ml.OJ.2 | 5585149 | 317824 | 24680 | 1134813 | 44057 | 78.6 |
| Ml.OJ.3 | 4709910 | 272067 | 21670 | 976167 | 47603 | 78.1 |
| Ml.OJ.4 | 4584617 | 266829 | 23372 | 880669 | 54025 | 78.9 |
| Ml.OJ.5 | 4891895 | 283361 | 23039 | 963523 | 49698 | 78.8 |
| Ml.PJ.1 | 5039454 | 267255 | 26124 | 1169256 | 49811 | 76.9 |
| Ml.PJ.2 | 5216451 | 278160 | 27837 | 1050162 | 54993 | 78.7 |
| Ml.PJ.3 | 3889142 | 224610 | 34629 | 727339 | 98504 | 78.2 |
| Ml.PJ.4 | 5251767 | 298476 | 29451 | 944191 | 70551 | 79.6 |
| Ml.PJ.5 | 4477837 | 258137 | 23336 | 833693 | 51929 | 79.3 |
| No.OJ.1 | 5395709 | 370500 | 26167 | 1186153 | 41410 | 76.9 |
| No.OJ.2 | 5619181 | 362078 | 32970 | 1078713 | 50007 | 78.7 |
| No.OJ.3 | 5823554 | 352085 | 28237 | 1214977 | 50064 | 78 |
| No.OJ.4 | 4473735 | 268227 | 22884 | 931205 | 40737 | 78 |
| No.OJ.5 | 6132995 | 396430 | 28533 | 1294114 | 41058 | 77.7 |
| No.PJ.1 | 5352739 | 339834 | 45726 | 1163403 | 115779 | 76.3 |
| No.PJ.2 | 5997956 | 342225 | 35373 | 907526 | 56537 | 81.7 |

| | | | | | | |
|---|---|---|---|---|---|---|
| No.PJ.3 | 4080045 | 229890 | 42161 | 789728 | 104422 | 77.8 |
| No.PJ.4 | 3945021 | 217824 | 35181 | 856809 | 71230 | 77 |
| No.PJ.5 | 3183942 | 234399 | 119507 | 816096 | 314206 | 68.2 |
| Pf.OJ.1 | 7947807 | 404248 | 29027 | 1394316 | 79816 | 80.6 |
| Pf.OJ.2 | 5791279 | 292900 | 29938 | 1334139 | 63852 | 77.1 |
| Pf.OJ.3 | 5161361 | 258603 | 24777 | 1126350 | 58339 | 77.9 |
| Pf.OJ.4 | 4563584 | 238081 | 25578 | 1330626 | 52181 | 73.5 |
| Pf.OJ.5 | 4791825 | 232390 | 26580 | 808562 | 65159 | 80.9 |
| Pf.PJ.1 | 7035279 | 345254 | 28508 | 1277919 | 84201 | 80.2 |
| Pf.PJ.2 | 3960377 | 201228 | 24972 | 849788 | 58079 | 77.7 |
| Pf.PJ.3 | 6174510 | 300868 | 38198 | 1298869 | 93285 | 78.1 |
| Pf.PJ.4 | 4795750 | 220939 | 19148 | 679422 | 46102 | 83.2 |
| Pf.PJ.5 | 5575886 | 263517 | 26392 | 813940 | 66816 | 82.6 |
| Pp.OJ.1 | 8314060 | 436690 | 25611 | 1611437 | 66925 | 79.5 |
| Pp.OJ.2 | 4829686 | 250440 | 18508 | 1107668 | 37463 | 77.4 |
| Pp.OJ.3 | 1519548 | 78655 | 13269 | 2173517 | 32834 | 39.8 |
| Pp.OJ.4 | 4737067 | 244597 | 69304 | 756769 | 170960 | 79.2 |
| Pp.OJ.5 | 5412469 | 298584 | 27446 | 935672 | 68760 | 80.3 |
| Pp.PJ.1 | 5315694 | 259600 | 15636 | 1123677 | 37141 | 78.7 |
| Pp.PJ.2 | 4682035 | 223223 | 17253 | 1075753 | 34994 | 77.6 |
| Pp.PJ.3 | 5074270 | 321073 | 212345 | 1344232 | 648179 | 66.8 |
| Pp.PJ.4 | 3516755 | 189054 | 130294 | 606269 | 298631 | 74.2 |
| Pp.PJ.5 | 4258398 | 210799 | 20458 | 626323 | 48020 | 82.5 |
| Prp.OJ.1 | 4714895 | 242922 | 31775 | 937027 | 50828 | 78.9 |
| Prp.OJ.2 | 6487462 | 363540 | 37729 | 1367218 | 89312 | 77.7 |
| Prp.OJ.3 | 6025899 | 337751 | 32587 | 1290339 | 63548 | 77.8 |
| Prp.OJ.4 | 7116572 | 405090 | 40104 | 1410959 | 96119 | 78.5 |
| Prp.OJ.5 | 7775110 | 409071 | 48075 | 1421183 | 114281 | 79.6 |
| Prp.PJ.1 | 5080689 | 249289 | 43275 | 1001970 | 83982 | 78.7 |
| Prp.PJ.2 | 4903357 | 249223 | 26562 | 1052028 | 55304 | 78 |
| Prp.PJ.3 | 5502725 | 288357 | 38907 | 1118220 | 92950 | 78.2 |
| Prp.PJ.4 | 3738386 | 200981 | 22490 | 704760 | 53341 | 79.2 |
| Prp.PJ.5 | 3305380 | 185694 | 18144 | 644620 | 43227 | 78.8 |
| Ps.OJ.1 | 6399897 | 372071 | 35014 | 1394940 | 69499 | 77.4 |
| Ps.OJ.2 | 5709661 | 311782 | 42946 | 1214005 | 82455 | 77.6 |
| Ps.OJ.3 | 4626788 | 251769 | 39102 | 871993 | 64375 | 79 |

| | | | | | | |
|---|---|---|---|---|---|---|
| Ps.OJ.4 | 5263877 | 280740 | 32167 | 975656 | 62177 | 79.6 |
| Ps.OJ.5 | 5181286 | 312443 | 30209 | 1056379 | 59093 | 78 |
| Ps.PJ.1 | 5740880 | 318404 | 43152 | 1209451 | 102270 | 77.4 |
| Ps.PJ.2 | 4352284 | 229640 | 54282 | 801223 | 96833 | 78.6 |
| Ps.PJ.3 | 6111323 | 324170 | 61472 | 1172280 | 135023 | 78.3 |
| Ps.PJ.4 | 5319457 | 267421 | 74648 | 947634 | 137690 | 78.8 |
| Ps.PJ.5 | 5048018 | 280401 | 52025 | 994187 | 114215 | 77.8 |
| Ptb.OJ.1 | 4563367 | 269206 | 18527 | 1063636 | 31511 | 76.7 |
| Ptb.OJ.2 | 7114957 | 419676 | 40571 | 1378785 | 81318 | 78.7 |
| Ptb.OJ.3 | 5302622 | 319323 | 25183 | 1098392 | 44805 | 78.1 |
| Ptb.OJ.4 | 5343635 | 318605 | 33327 | 1028340 | 67922 | 78.7 |
| Ptb.OJ.5 | 5970463 | 357750 | 33511 | 1165518 | 64540 | 78.6 |
| Ptb.PJ.1 | 3742805 | 212035 | 17935 | 852540 | 36475 | 77 |
| Ptb.PJ.2 | 4834135 | 277458 | 31463 | 838624 | 73872 | 79.8 |
| Ptb.PJ.3 | 4238648 | 252073 | 19920 | 939627 | 37871 | 77.2 |
| Ptb.PJ.4 | 4388179 | 247138 | 31849 | 842065 | 69195 | 78.7 |
| Ptb.PJ.5 | 4644086 | 269247 | 34640 | 766917 | 79356 | 80.1 |
| Py.OJ.1 | 5347906 | 295047 | 25753 | 958661 | 52108 | 80.1 |
| Py.OJ.2 | 6205819 | 349638 | 25199 | 1147214 | 45857 | 79.8 |
| Py.OJ.3 | 5661376 | 306552 | 24603 | 1038562 | 52388 | 79.9 |
| Py.OJ.4 | 4897704 | 263633 | 23936 | 938567 | 49366 | 79.3 |
| Py.OJ.5 | 4524067 | 246387 | 25532 | 796194 | 61593 | 80 |
| Py.PJ.1 | 5602097 | 301933 | 24513 | 942736 | 45650 | 81 |
| Py.PJ.2 | 6906283 | 379801 | 28108 | 1130821 | 50976 | 81.3 |
| Py.PJ.3 | 5609316 | 301372 | 25567 | 938159 | 54045 | 81 |
| Py.PJ.4 | 3640462 | 194391 | 19267 | 552975 | 40483 | 81.9 |
| Py.PJ.5 | 4516777 | 246730 | 23451 | 736712 | 47925 | 81.1 |
| Sd.OJ.1 | 5539180 | 299107 | 34642 | 1048749 | 86928 | 79 |
| Sd.OJ.2 | 5783309 | 286283 | 26056 | 1047359 | 56147 | 80.3 |
| Sd.OJ.3 | 4732049 | 229358 | 17728 | 1062104 | 35849 | 77.9 |
| Sd.OJ.4 | 4975049 | 251903 | 22510 | 833429 | 51552 | 81.1 |
| Sd.OJ.5 | 5941520 | 294972 | 25440 | 1001098 | 65225 | 81.1 |
| Sd.PJ.1 | 3772485 | 209931 | 51991 | 879914 | 113635 | 75 |
| Sd.PJ.2 | 7119169 | 347200 | 62171 | 1455446 | 158161 | 77.9 |
| Sd.PJ.3 | 6914250 | 328658 | 59628 | 1553180 | 144880 | 76.8 |
| Sd.PJ.4 | 5696568 | 261873 | 25096 | 779123 | 53850 | 83.6 |

| | | | | | | |
|---|---|---|---|---|---|---|
| Sd.PJ.5 | 5086543 | 221518 | 23637 | 724132 | 53164 | 83.3 |
| Sf.OJ.1 | 7143170 | 433689 | 30287 | 1463953 | 52903 | 78.3 |
| Sf.OJ.2 | 7146958 | 448993 | 35039 | 1358082 | 58811 | 79 |
| Sf.OJ.3 | 5516493 | 357074 | 27027 | 1148832 | 46124 | 77.7 |
| Sf.OJ.4 | 6186658 | 386130 | 26044 | 1268508 | 40333 | 78.2 |
| Sf.OJ.5 | 5820808 | 308921 | 30235 | 960451 | 62498 | 81 |
| Sf.PJ.1 | 3776798 | 220861 | 14725 | 835601 | 25838 | 77.5 |
| Sf.PJ.2 | 6219309 | 396525 | 33331 | 1162388 | 67773 | 78.9 |
| Sf.PJ.3 | 4308855 | 261284 | 23294 | 804396 | 43510 | 79.2 |
| Sf.PJ.4 | 5041048 | 316753 | 24023 | 955537 | 43376 | 79 |
| Sf.PJ.5 | 4924204 | 275088 | 56812 | 858274 | 149728 | 78.6 |
| Tm.OJ.1 | 8516062 | 434545 | 30617 | 1385654 | 66778 | 81.6 |
| Tm.OJ.2 | 5596680 | 296955 | 27078 | 1244071 | 55597 | 77.5 |
| Tm.OJ.3 | 5345382 | 280030 | 38934 | 1041742 | 85567 | 78.7 |
| Tm.OJ.4 | 3757697 | 201681 | 31147 | 806765 | 67636 | 77.2 |
| Tm.OJ.5 | 4229479 | 225236 | 33997 | 633138 | 87473 | 81.2 |
| Tm.PJ.1 | 7627311 | 405288 | 27440 | 1403580 | 64795 | 80 |
| Tm.PJ.2 | 4940304 | 263708 | 21555 | 979902 | 47495 | 79 |
| Tm.PJ.3 | 3027989 | 214018 | 144718 | 767184 | 494718 | 65.1 |
| Tm.PJ.4 | 3788748 | 212760 | 40825 | 789187 | 132028 | 76.3 |
| Tm.PJ.5 | 3703931 | 209783 | 62288 | 533882 | 138465 | 79.7 |
| Tt.OJ.1 | 5390289 | 291650 | 18175 | 1151114 | 36510 | 78.3 |
| Tt.OJ.2 | 4700444 | 274178 | 19728 | 941133 | 33420 | 78.7 |
| Tt.OJ.3 | 6832043 | 393398 | 27203 | 1332545 | 45741 | 79.2 |
| Tt.OJ.4 | 3735283 | 219732 | 24905 | 704799 | 53997 | 78.8 |
| Tt.OJ.5 | 5699037 | 323644 | 34887 | 998449 | 91483 | 79.7 |
| Tt.PJ.1 | 4001285 | 239133 | 14062 | 873044 | 24723 | 77.7 |
| Tt.PJ.2 | 4129526 | 243516 | 14865 | 968433 | 24749 | 76.7 |
| Tt.PJ.3 | 5274110 | 293746 | 21493 | 979832 | 44297 | 79.7 |
| Tt.PJ.4 | 4472595 | 269953 | 30501 | 816241 | 68444 | 79.1 |
| Tt.PJ.5 | 5180742 | 288256 | 24341 | 979550 | 55508 | 79.4 |

\

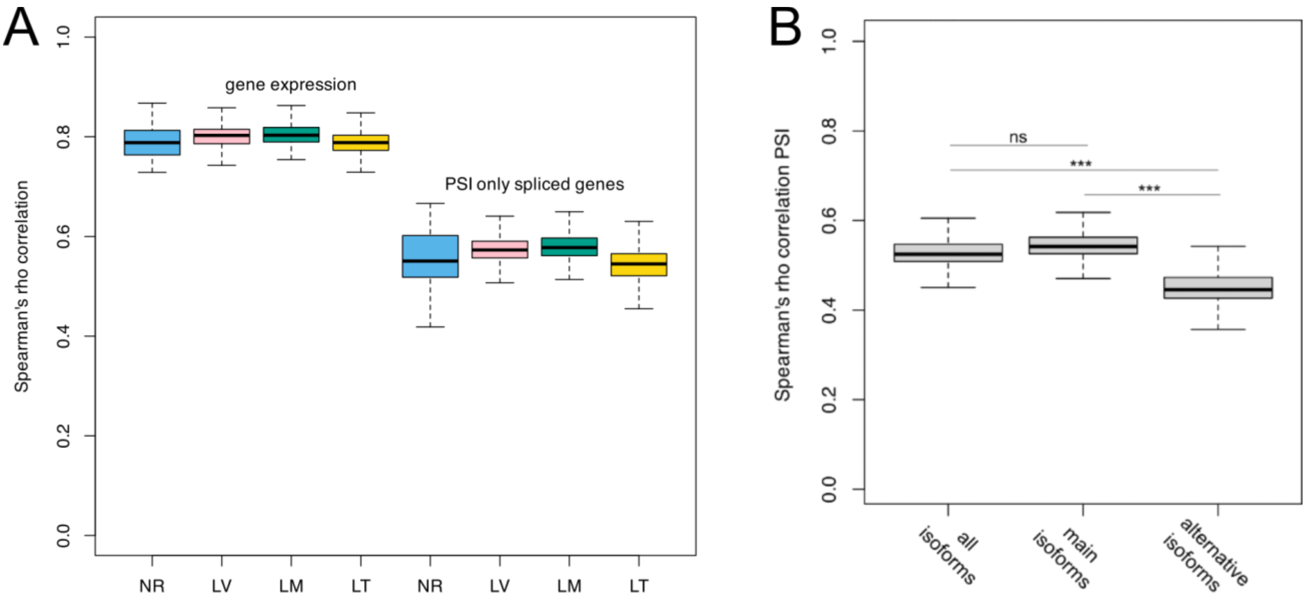

**Figure S1 Spearman's rank correlation coefficient (ρ) of gene expression (GE) and percent spliced in (PSI)**

(A) Boxplots of Spearman's ρ of PSI and GE calculated per lake (LT: Lake Tanganyika; LM:Lake Malawi; LV: Lake Victoria) as well as non-radiating species (NR). PSI is only calculated for genes with more than one isoform. All within lake and within non-radiating comparisons of gene expression rho versus PSI rho are significantly different (two-sided t-test $p < 0.001$). (B) Conservation of main and alternative isoforms in cichlid adaptive radiations. Distribution of Spearman's rank correlation coefficient (ρ) of isoform percent spliced in (PSI) between pairs of samples (n=200) for all genes with more than one isoform. Isoforms are categorised into main (constitutive) isoforms and alternative isoforms. Significant difference is calculated using two-sided t-test (*** = $p < 0.001$, ns = not significant).

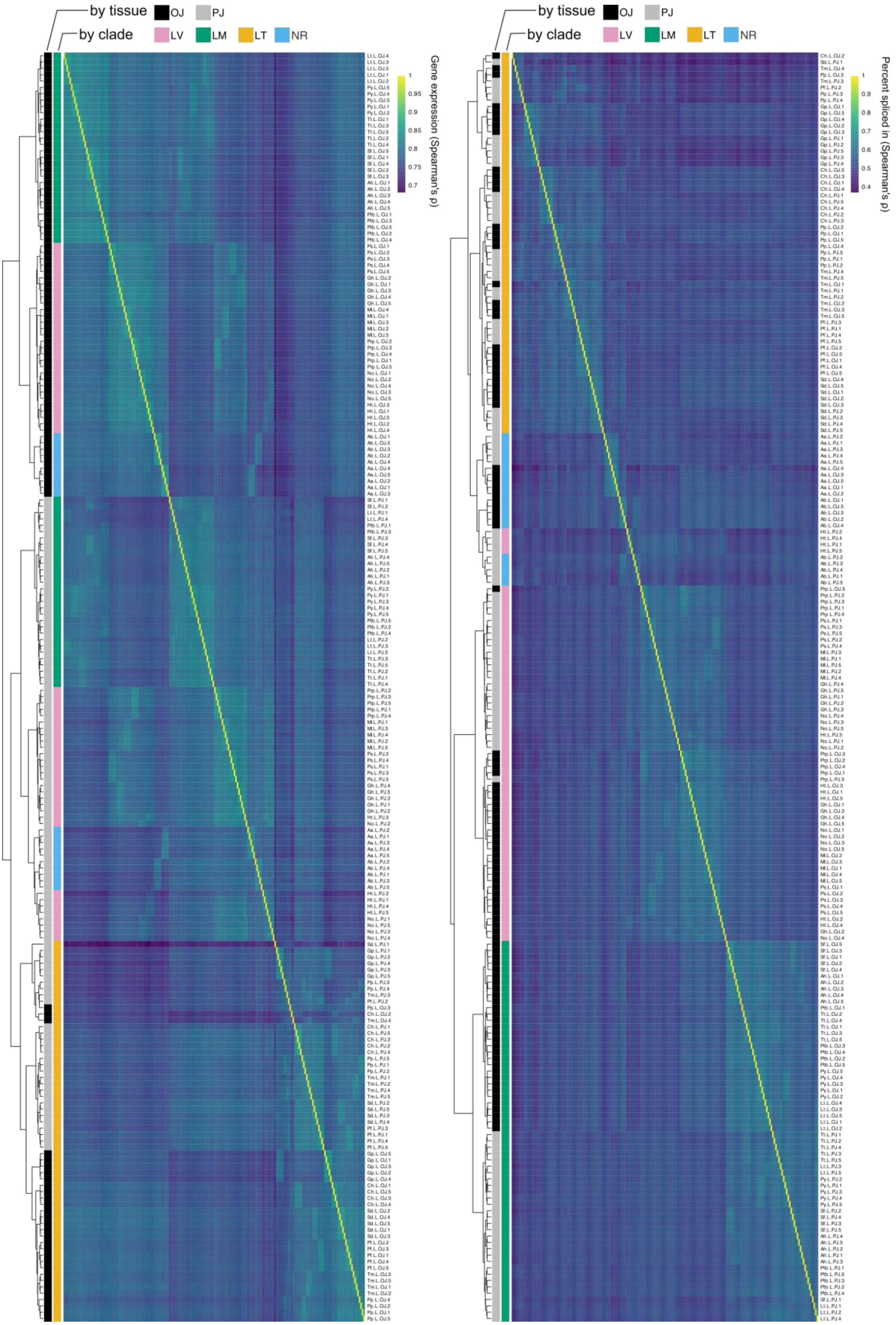

by tissue
OJ
PJ
by clade
LV
LM
LT
NR
Gene expression (Spearman's ρ)
1
0.95
0.9
0.85
0.8
0.75
0.7
by tissue
OJ
PJ
by clade
LV
LM
LT
NR
Percent spliced in (Spearman's ρ)
1
0.9
0.8
0.7
0.6
0.5
0.4

**Figure S2 Profiling of alternative splicing and gene expression in cichlid radiations.**

The same as main Figure 2 but with species names and tissue annotated next to rows. Hierarchical clustering of Spearman's rank correlation coefficient (ρ) of normalised gene expression and isoform percent spliced in (PSI) between pairs of samples (n=200). Sample clustering is represented as a tree on the left side of the heatmaps, with tissue and clade (radiations or nonradiating) of origin of species annotated in colours. *Paralabidochromis sauvagei* (Ps), *Mbipia lutea* (Ml), *Neochromis omnicaeruleus* (No), *Gaurochromis hiatus* (Gh), *Prognathochormis perrieri* (Pp), *Haplochromis thereuterion* (Ht), *Aulonocara hansbaenschi* (Ah), *Labeotropheus trewavasae* (Lt), *Petrotilapia* sp. 'Thick bars' (Ptb), *Petrotilapia* sp. 'Yellow chin Chewere' (Py), *Sciaenochromis fryeri* (Sf), *Tropheops tropheops* (Tt), *Ctenochromis horei* (Ch), *Gnathochromis pfefferi* (Gp), *Petrochromis famula* (Pf), *Petrochromis polyodon* (Pp), *Simochromis diagramma* (Sd), *Tropheus moorii* (Tm), *Astatoreochromis alluaudi* (Aa), *Astatotilapia burtoni* (Ab).

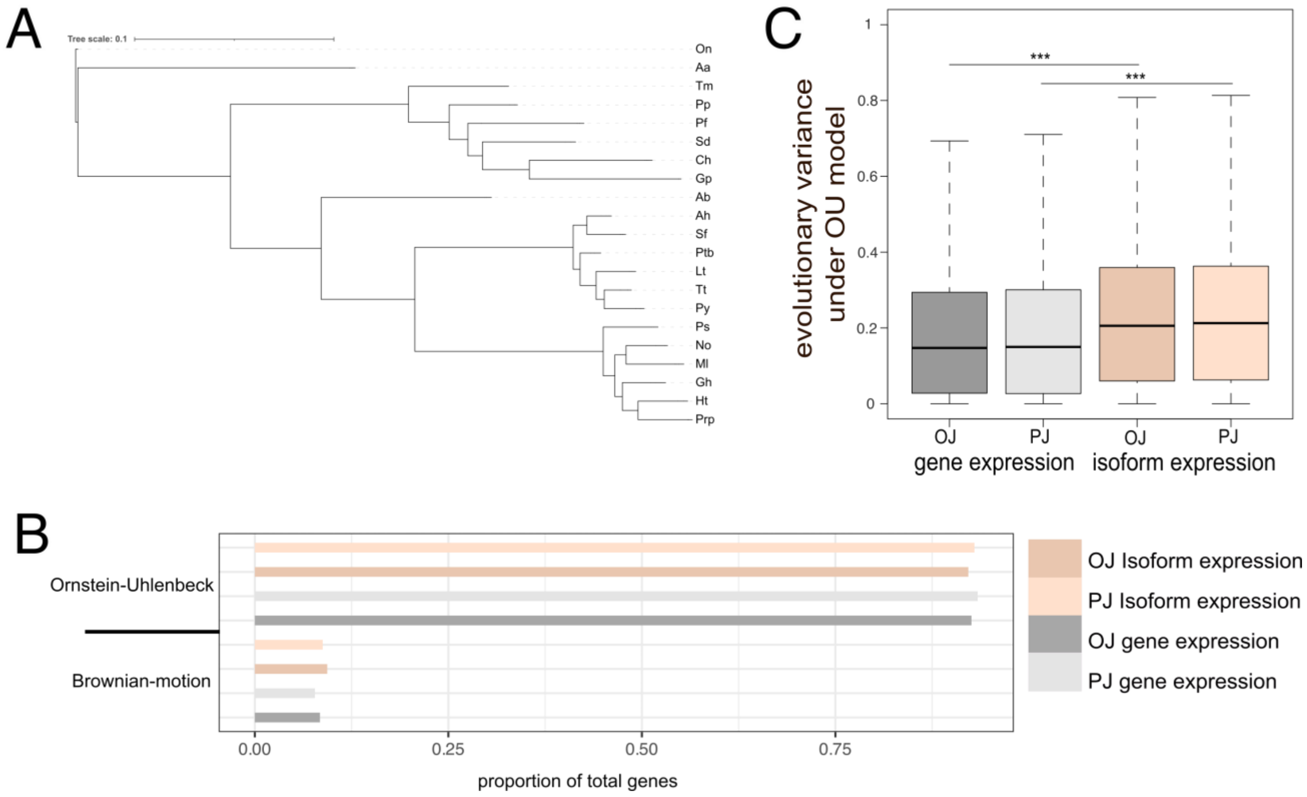

**Figure S3 Evolution of transcriptome-wide gene and isoform expression in cichlid radiations.**

(A) Phylogenetic reconstruction of all species in this study using transcriptomic SNPs. (B) Evolutionary variance in gene and isoform expression along the phylogeny under the Ornstein-Uhlenbeck (OU) model. (C) Number of genes and isoforms whose expression evolution along the phylogeny fits the Brownian motion (BM) or Ornstein-Uhlenbeck (OU) models.

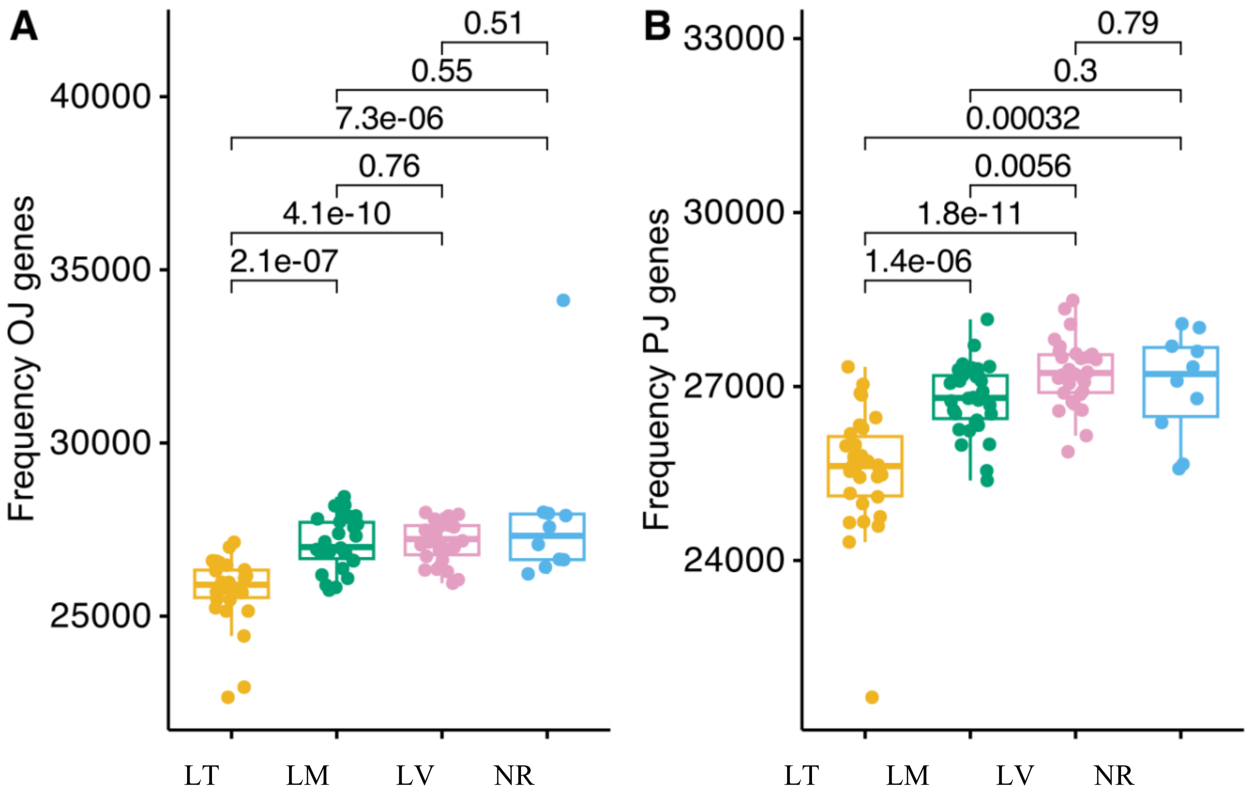

**Figure S4 Transcriptomic variation in non-radiating and radiating cichlids: genes expressed.** (A, B) The absolute number of expressed genes identified in the oral and pharyngeal jaw mRNA of all samples. *P*-values from Mann-Whitney U test between groups is shown. Groups: Lake Victoria (LV), Lake Malawi (LM) and Lake Tanganyika (LT) compared to non-radiating (NR)

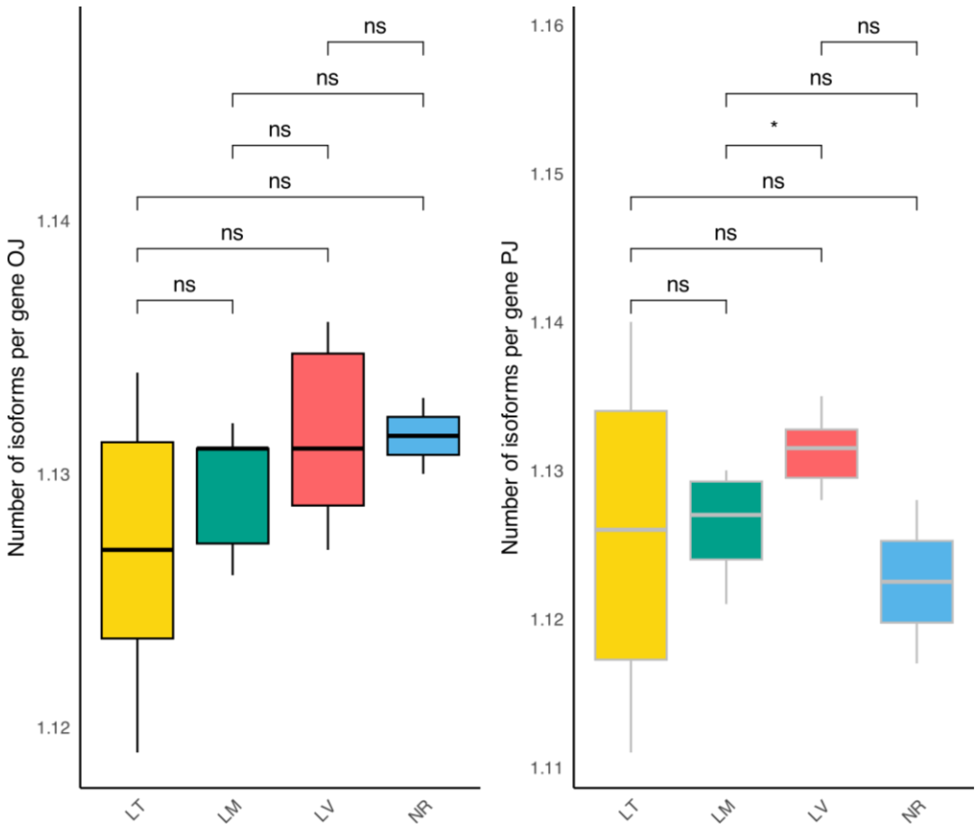

**Figure S5 Transcriptomic variation in non-radiating and radiating cichlids: number of isoforms per gene expressed.**

(A, B) The absolute number of expressed genes identified in the oral and pharyngeal jaw mRNA of all samples. Asterisks denote Mann-Whitney U test significance: $p < 0.05$* and NS: no significance. Groups: Lake Victoria (LV), Lake Malawi (LM) and Lake Tanganyika (LT) compared to non-radiating (NR)

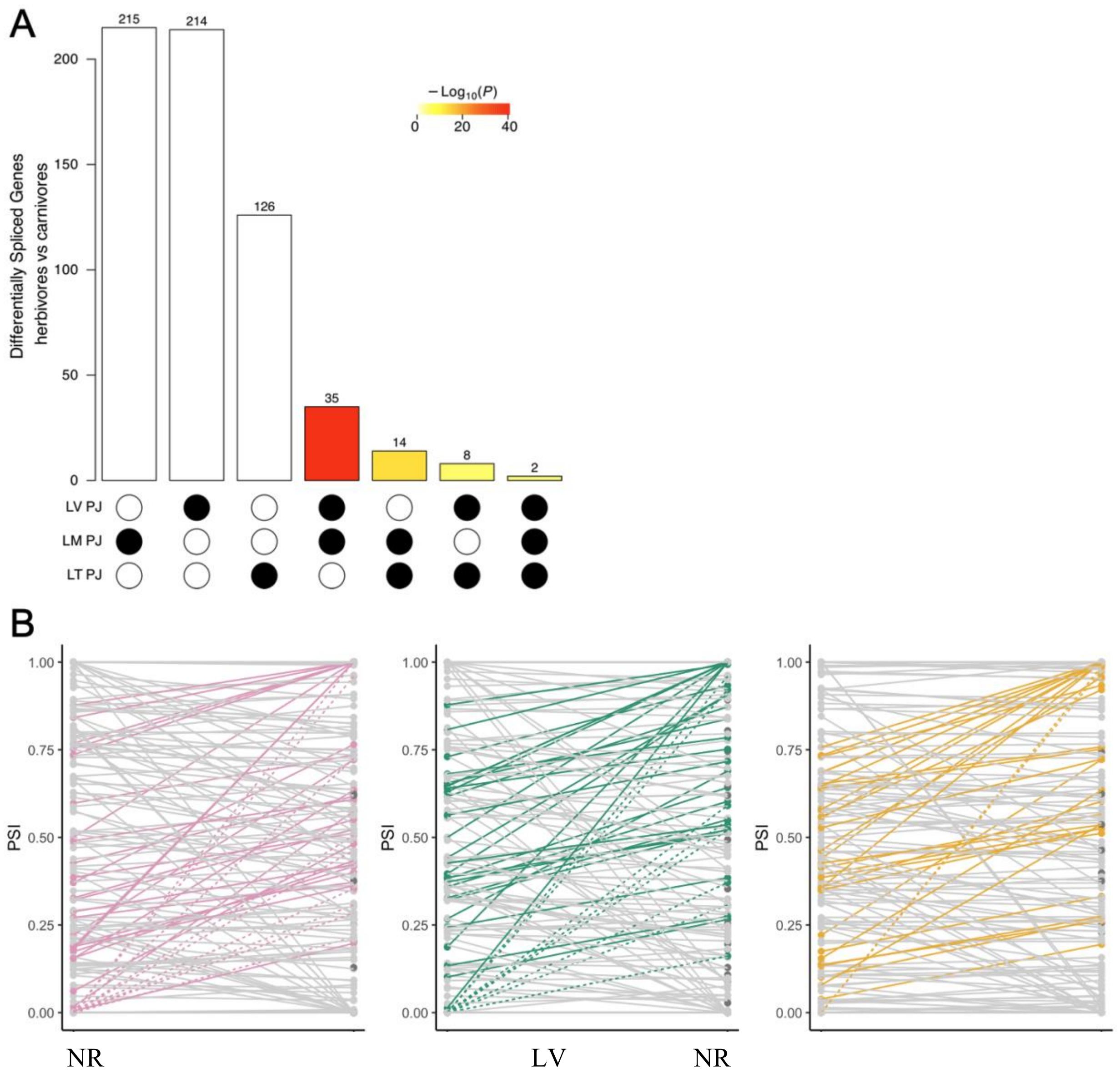

**Figure S6 Evolutionary dynamics of differentially spliced genes between pharyngeal jaws (PJ) of herbivores and carnivores from each radiation.**

(A) Number of significantly differentially spliced genes ($q < 0.05$) in PJ's of carnivores vs herbivores with significant overlaps shown (hypergeometric test *p* values are illustrated as heatmap). (B) Testing the hypothesis depicted in main Figure 3D in our transcriptome data from Lake Victoria (LV), Lake Malawi (LM) and Lake Tanganyika (LT) compared to non-radiating (NR) species that share a common ancestor with these radiations. Testing the hypothesis depicted in (D) in our transcriptome data from Lake Victoria radiation (LV), Lake Malawi radiation (LM) and Lake Tanganyika radiation (LT) compared to non-radiating species (NR). Mean PSI was calculated all individuals of all species in each group (NR, LV, LM, LT). All lines represent all isoforms of genes that were significantly differentially spliced between herbivores and carnivores within each radiation. Solid grey lines represent all isoforms that were expressed at lower or similar levels in the NR species and species from each radiation (PSI differences <= 0.2). Dotted grey lines represent all isoforms that were not expressed in NR species but were expressed species from each radiation (PSI differences <= 0.2). Coloured solid lines indicate isoforms that were expressed at low levels in NR species but were higher expressed in radiating species (PSI differences > 0.2). Coloured dotted lines indicate isoforms that were not expressed in NR species but were expressed in radiating species (PSI differences > 0.2).

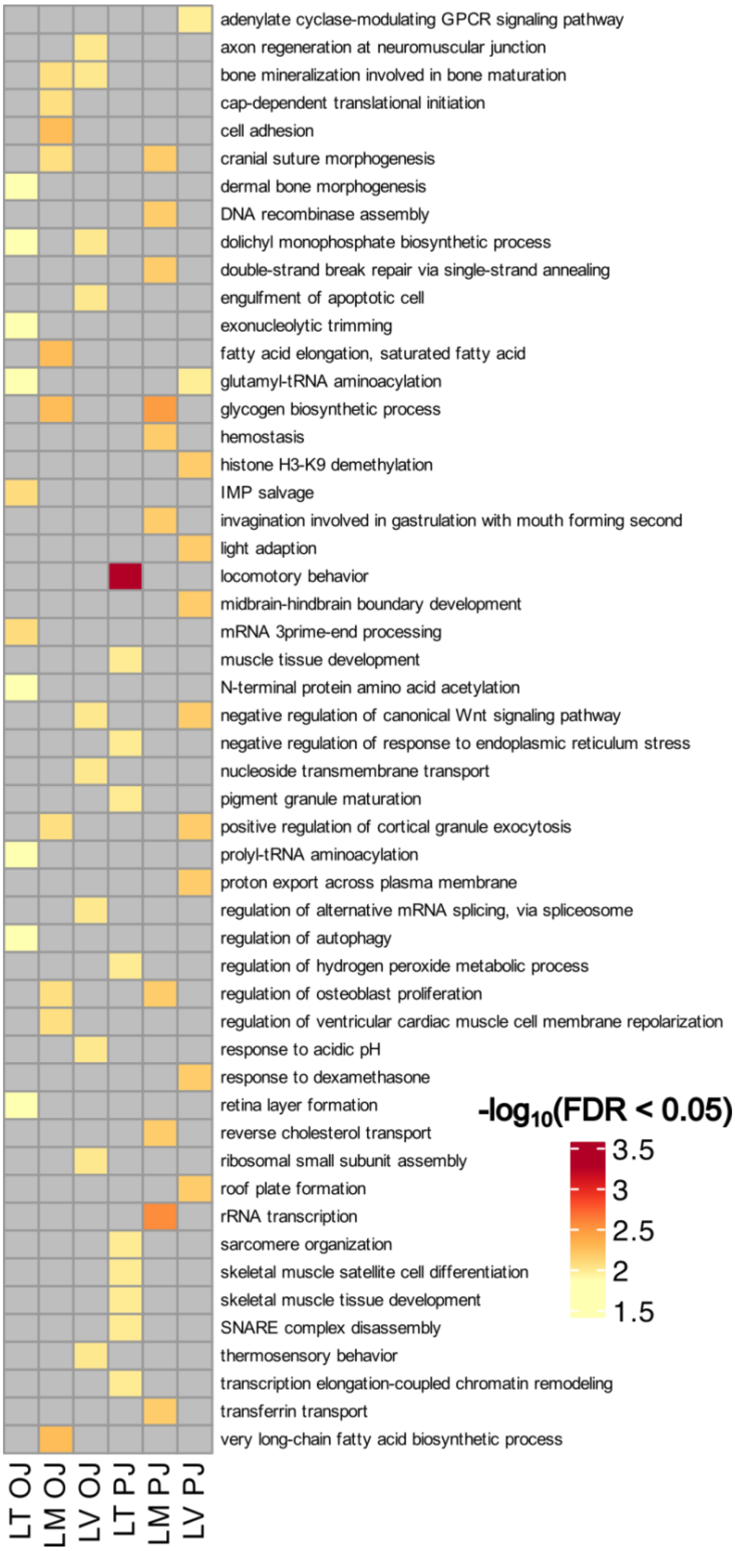

**Figure S7 Gene Ontology enrichment of differentially spliced genes between herbivores and carnivores for oral (OJ) and pharyngeal jaws (PJ).**

Only significant terms shown after correction for multiple testing (Fisher's exact test $q < 0.05$). LV: Lake Victoria, LM: Lake Malawi, LT: Lake Tanganyika, NR: non-radiating.

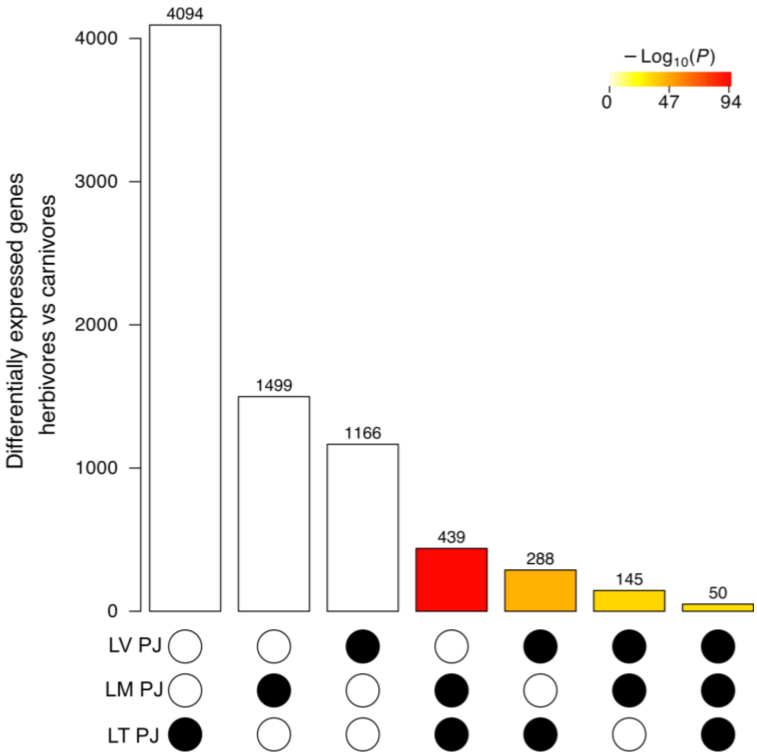

**Figure S8 Gene expression divergence in pharyngeal jaws of herbivore and carnivore species.**

Number of differentially expressed genes (DEGs, $q < 0.05$) between pharyngeal jaws (PJ) of herbivores and carnivores within Lake Victoria (LV), Lake Malawi (LM) and Lake Tanganyika (LT), with significantly greater overlaps than expected by chance are shown as a colour heatmap (hypergeometric test $p$ values).

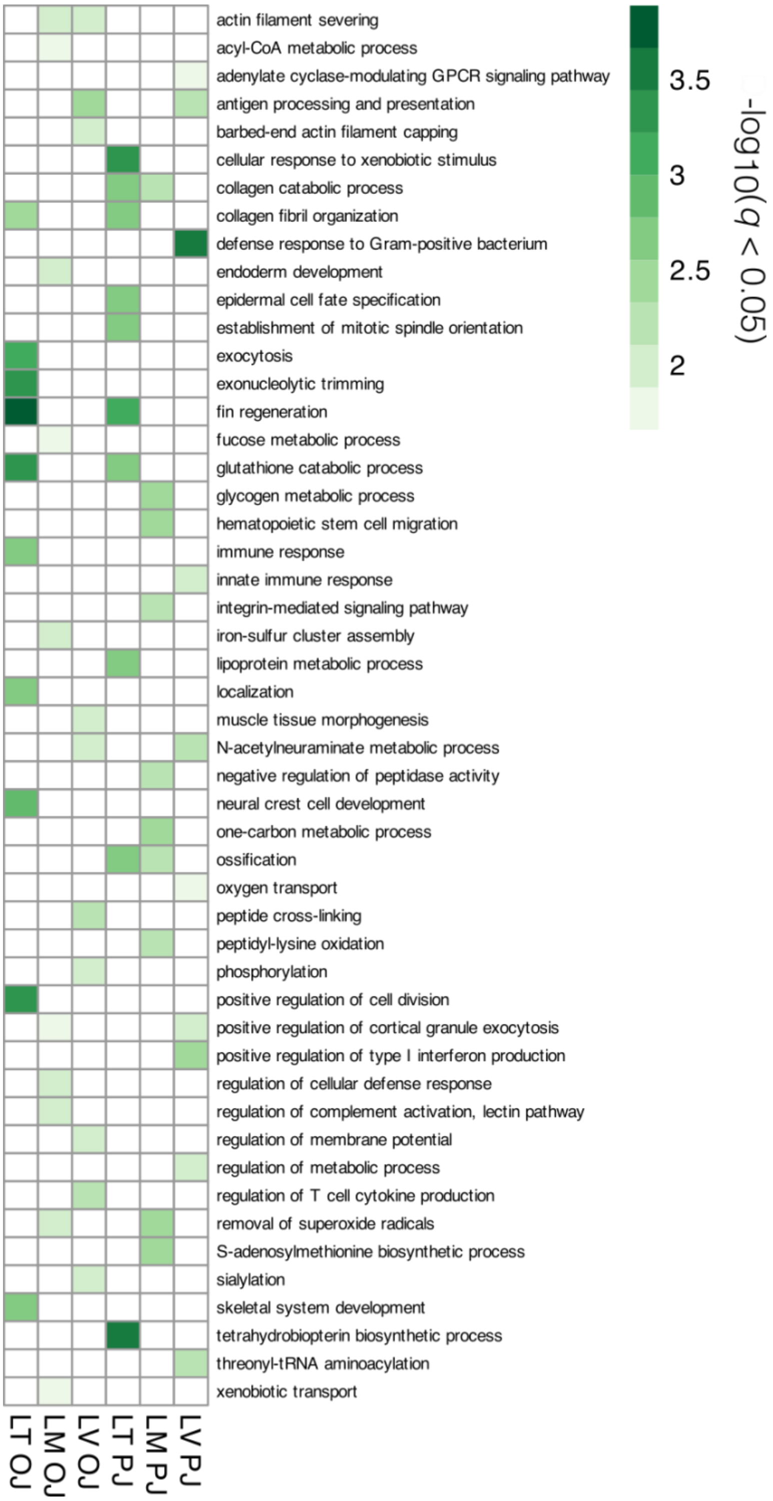

**Figure S9 Gene Ontology enrichment of differentially expressed genes between herbivores and carnivores.**

Only significant terms shown after correction for multiple testing (Fisher's exact test $q < 0.05$).

**Supplementary Materials and Methods: qPCR analysis**

From 4 replicates of RNA samples of each jaw, stage and species, we used 500ng of RNA to synthesize first strand cDNA using High Capacity cDNA Reverse Transcription kit (Applied Biosystems). The cDNA from each sample was diluted 1:10 times in nuclease-free water to be used for qPCR. We used *rpl18* as a reference gene, which is validated as a stably expressed gene in both cichlid jaws across different trophic niches(1) to normalize our expression data for qPCR. The primers were designed at conserved sequence regions of the coding sequences from all species using CLC Genomic Workbench (v7.5 CLC Bio, Aarhus, Denmark) for alignment and delineation of exon boundaries using *A. burtoni* genome(2). Primer Express 3.0 (Applied Biosystems, CA, USA) and OligoAnalyzer 3.1 (Integrated DNA Technology) were used to design the primers as described in(1, 3). Maxima SYBR Green/ROX qPCR Master Mix (2X) (Thermo Fisher Scientific, Germany) was used to generate qPCR reactions. The amplification steps were conducted in 96 well-PCR plates on ABI 7500 real-time PCR System (Applied Biosystems). The primer efficiency calculation, qPCR program, plate set-up and a dissociation step were performed as described in(1, 3). To normalize the expression levels, the Cq value of *rpl18* was used as normalization factor (Cq reference), and ΔCq of each target gene was calculated accordingly (ΔCq target = Cq target – Cq reference). Relative expression quantities (RQ) were calculated for the normalized values using $E^{-\Delta\Delta Cq}$ and then fold difference values were calculated by transformation of RQ values to logarithmic values in order to conduct further statistical analysis. The statistical expression differences were identified using ANOVA statistical tests, followed by Tukey's HSD post hoc tests.

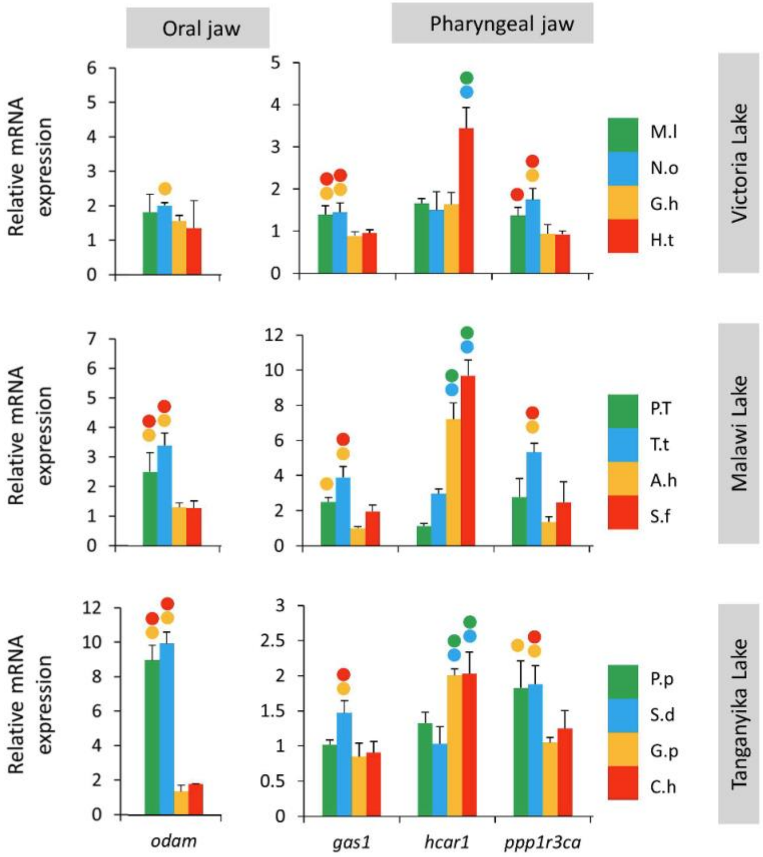

**Figure S10 quantitative PCR (qPCR) validation of convergently expressed genes between herbivores and carnivores in all three lakes.**

Blue/Green = herbivore species. Orange/Red – carnivore species. Species names are abbreviated. See Table 1 for full names. Filled circles above bars depict significantly different comparisons among groups.

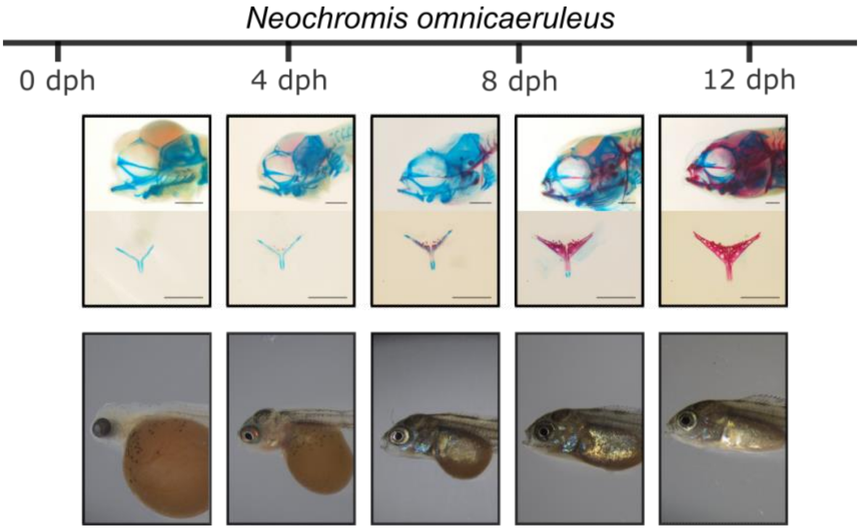

**Figure S11 Development of external morphology and craniofacial bone and cartilage in *Neochromis omnicaeruleus* from hatching to developmental stage 26 (end of postembryonic development).**

Top row: Alizarin red (bone) and Alcian blue (cartilage) stained images of *N. omnicaeruleus* sampled at regular intervals from the hatching to 12 days post hatching (dph). Images provided by B Dreo. Bottom row: external morphology of *N. omnicaeruleus* sampled at regular intervals over 12 dph, when egg yolk is absorbed into the body cavity and which signifies the end of the larval stage and the beginning of the juvenile stage, ad individuals leave the protection of the mother's buccal cavity to begin their independent life and feeding. Images provided by C Mlay.

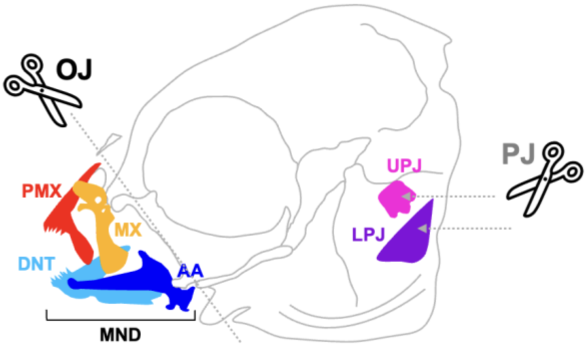

**Figure S12 Dissection strategy of oral and pharyngeal jaws (OJ, PJ) of stage 26 larvae used for RNA extractions.**

Oral jaw dissection includes the following bones: angulo-articular (AA), dentary (DNT), mandible (MND), maxilla (MX), pre-maxilla (PMX), as well as surrounding cartilage and muscle. Pharyngeal jaw dissection includes the upper pharyngeal jaw (UPJ) and the lower pharyngeal jaw (LPJ) as well as surrounding cartilage and muscle. For both OJ and PJ, we cleaned off as much surrounding tissue as possible under a stereomicroscope.

**Supplementary Materials and Methods: Geometric morphometric analysis**

We scanned the mandibles of 17 out of the 18 lacustrine study species, each represented by three adult male individuals, resulting in a total of 51 specimens analysed. According to our study design we bred and raised all specimens in our aquarium facility at the University of Graz under a standardised rearing and diet regime. We excluded the two riverine outgroups *Astatotilapia burtoni* and *Astatoreochromis alluaudi*, as well as the Lake Victoria endemic *Prognathochromis perrieri* for which only wild-caught specimens were available. The study species were classified in three trophic niches: omnivore, herbivore and carnivore (see supporting information Table S3). We used the SCANCO MEDICAL µCT 40 scanner to obtain the computer tomography scans and further reconstructed three-dimensional volumes by using the analytic software 3D Slicer (4). As studies in cichlids have shown that the craniofacial region comprises a major source of interspecific variation (5, 6), we focused on the mandible of the lower oral jaw. We defined a three-dimensional landmark set twelve homologous points which were uniformly well detectable on all study species (Fig. S13). Due to the oral symmetry of the used species, the left side was deemed sufficient for the morphometric analysis. For the descriptive termini of the precise landmark positions, we followed the classic paper (7). The complete landmark set is shown in Fig. S13 A to C, from different angles. For better visualization of the landmark positions a video is linked in the supporting information Video S1.

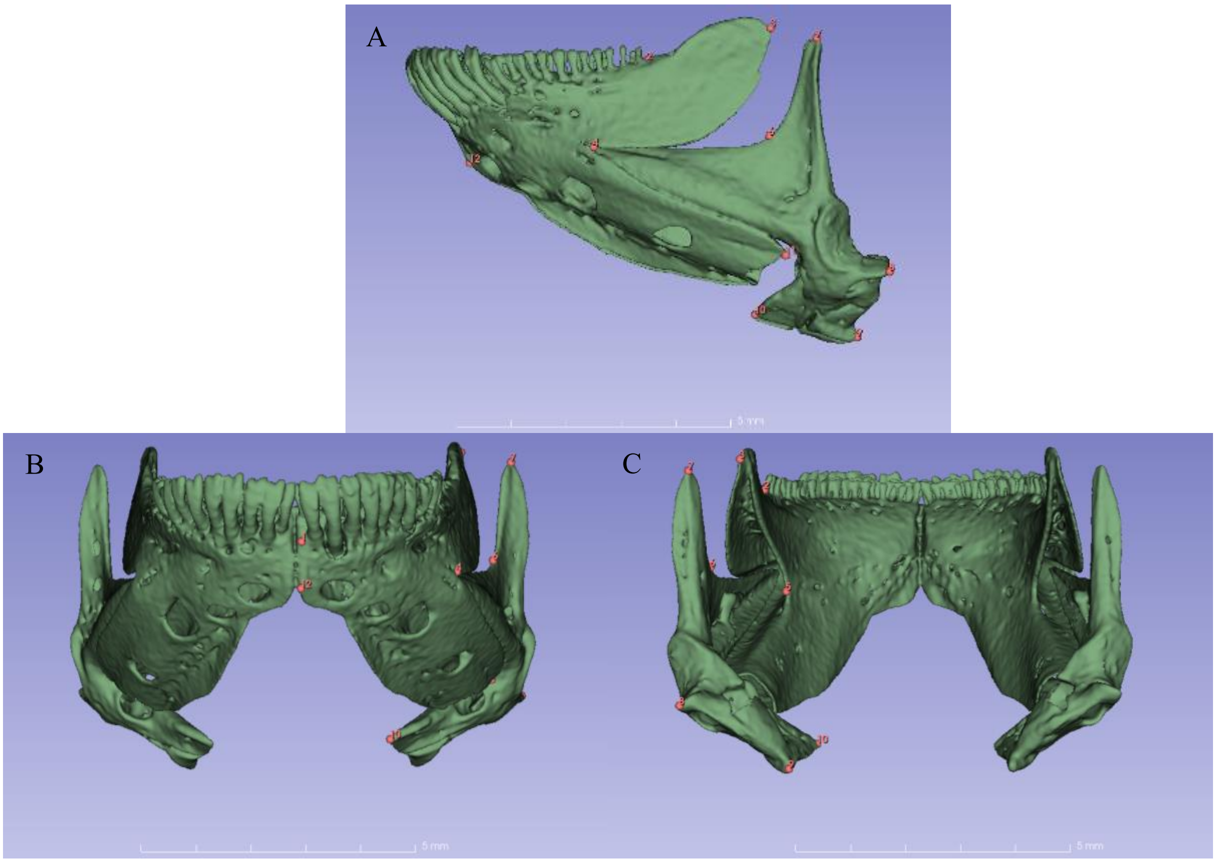

**Figure S13 Landmark template shown on a specimen of the Lake Victoria endemic *Neochromis omnicaeruleus*.**

(A) lateral view of the left side of the mandible. (B) anterior view. (C) posterior view. Number and position of landmarks: 1 - anterior dorsal teeth base, 2 - posterior teeth base, 3 - posterior tip of coronoid process, 4 - anterior tip of reentrant angle, 5 - dental fossa basin, 6 - articular web basin, 7 - post articular process tip, 8 - post articular process tip, 9 - posterior tip retroarticular process, 10 - anterior tip retroarticular process, 11 - posterior tip of ventral crest, 12 - anterior ventral tip of the dentary. The scale bar is 5 millimetres wide.

For the digitization of all 51 reconstructed mandible volumes and the generalized procrustes analysis (GPA) we used the integrated modules of 3D Slicer. We used the software package Past (8). We performed a PCA of all specimens (Fig. S14) and visualized shape differences between species with deformation grids of three-dimensional mandible volumes (Fig, S15) using 3D Slicer.

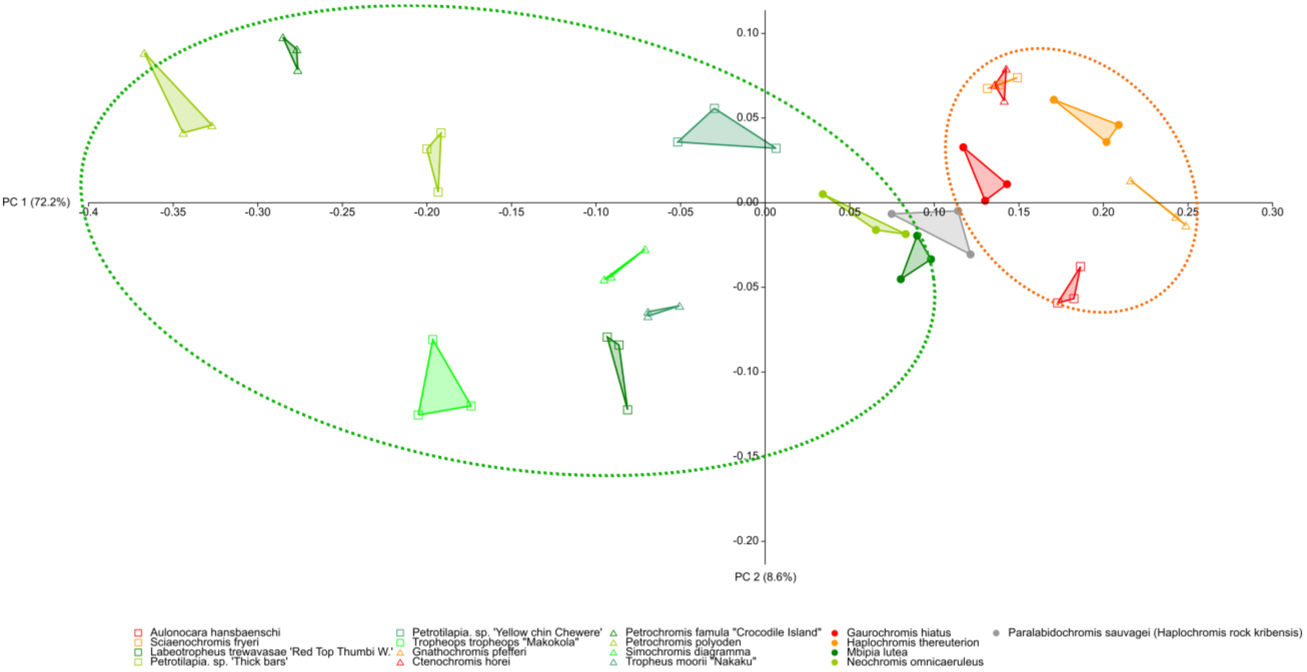

**Figure S14 Principal Component analysis (PCA) of 17 species from our studyplotted with lake-specific symbols (square LM; triangle LT; circle LV) and colour codes for trophic niches (carnivore red and orange; herbivore greenish colours and the omnivore in grey).**

B

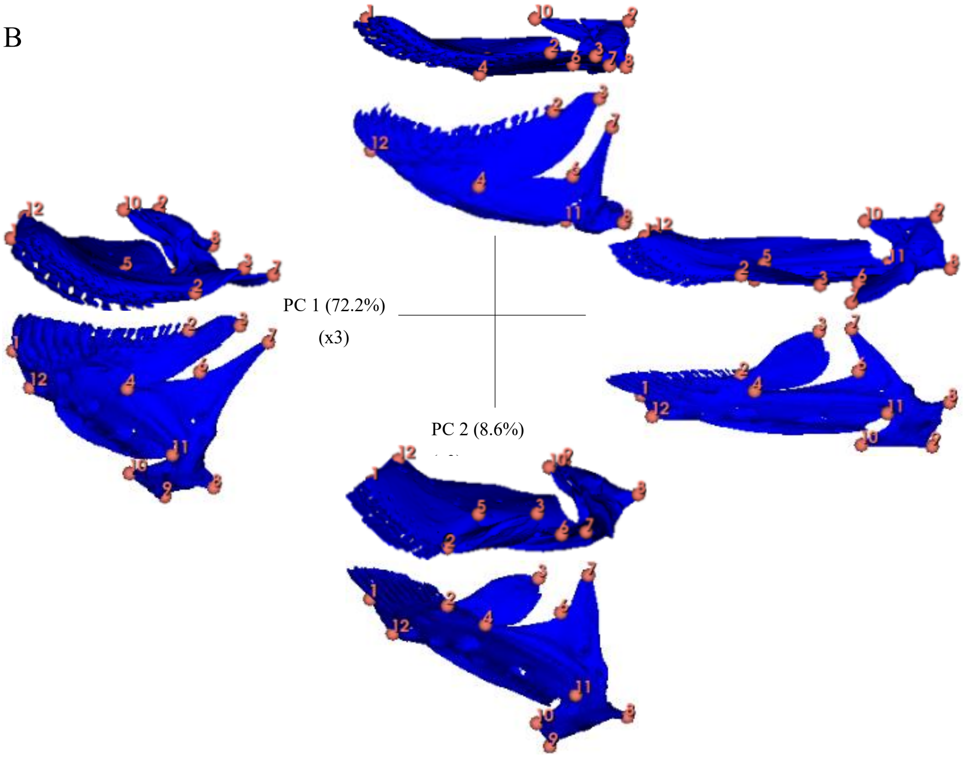

**Figure S15 Negative and positive deformation of extrapolated mandible half PC1(x-axis) and PC2 (y-axis) at dorsal view (top) and lateral view (bottom). All deformation grids were extrapolated by the factor three.**

**Supplementary Results and Discussion: Geometric morphometric analysis**

The first two principal components of the PCA explained total 80.8% of the variance: PC1 explained 72.2% of the variance, PC2 8.9% of the variance (Fig. S14). The separation of species regarding their trophic niche along the PC1-axies is distinctly visible for all lakes, with the herbivores clustering on the left side and the carnivore species clustering on the right side of the PC1 morphospace, with the omnivore *Paralabidochromis sauvagei* as intermediate. Within Lake Victoria the trend of trophic niche diverging along PC1 is visible, in line with the fact that Lake Victoria represents the youngest cichlid radiation between the three Great African Lakes. Lake Tanganyika as the oldest radiation shows the widest cluster distances among its species, and the Lake Malawi species comprise the second largest cluster range. Thus, Lake Victoria species show less interspecific mandible shape variation compared to Lakes Tanganika and Malawi, but species with similar trophic adaptations (herbivory vs carnivory) from each lake are diverging along parallel axes, consistent with previous findings (9).

The negative extrapolation along the PC1, shows a compressed mandible volume shape typical of herbivores (Fig. S15). In comparison the more elongated mandible volume along the positive extrapolation of PC1, similar to the shape of carnivores. Along the PC2-axis no separation between the species is possible, but the skewing of the articular process (Landmark 8 to Lankmark 10) is visible here.

**Table S3: Sampling list for geometric morphometric analyses.**

| lake | diet | species | nr. | ID |
|---|---|---|---|---|
| LT | carnivore | *Gnathochromis pfefferi* | Gnpf_A_01 | 543 |
| LT | carnivore | *Gnathochromis pfefferi* | Gnpf_A_02 | 544 |
| LT | carnivore | *Gnathochromis pfefferi* | Gnpf_A_03 | 528 |
| LT | herbivore | *Tropheus moorii "Nakaku"* | Trmo_A_01 | 460 |
| LT | herbivore | *Tropheus moorii "Nakaku"* | Trmo_A_02 | 461 |
| LT | herbivore | *Tropheus moorii "Nakaku"* | Trmo_A_03 | 462 |
| LT | herbivore | *Simochromis diagramma* | Sidi_A_01 | 247 |
| LT | herbivore | *Simochromis diagramma* | Sidi_A_02 | 254 |
| LT | herbivore | *Simochromis diagramma* | Sidi_A_03 | 255 |
| LT | herbivore | *Petrochromis famula "Crocodile Island"* | Pefa_A_01 | 426 |
| LT | herbivore | *Petrochromis famula "Crocodile Island"* | Pefa_A_02 | 441 |
| LT | herbivore | *Petrochromis famula "Crocodile Island"* | Pefa_A_03 | 442 |
| LT | herbivore | *Petrochromis polyodon* | Pepo_A_01 | 469 |
| LT | herbivore | *Petrochromis polyodon* | Pepo_A_02 | 534 |
| LT | herbivore | *Petrochromis polyodon* | Pepo_A_03 | 538 |
| LT | carnivore | *Ctenochromis horei* | Ctho_A_01 | 393 |
| LT | carnivore | *Ctenochromis horei* | Ctho_A_02 | 394 |
| LT | carnivore | *Ctenochromis horei* | Ctho_A_03 | 395 |
| LV | omnivore | *Paralabidochromis sauvagei (Haplochromis rock kribensis)* | Pasa_A_01 | 451 |
| LV | omnivore | *Paralabidochromis sauvagei (Haplochromis rock kribensis)* | Pasa_A_02 | 521 |
| LV | omnivore | *Paralabidochromis sauvagei (Haplochromis rock kribensis)* | Pasa_A_03 | 522 |
| LV | herbivore | *Mbipia lutea* | Mblu_A_01 | 525 |
| LV | herbivore | *Mbipia lutea* | Mblu_A_02 | 526 |
| LV | herbivore | *Mbipia lutea* | Mblu_A_03 | 527 |
| LV | carnivore | *Gaurochromis hiatus* | Gahi_A_01 | 483 |
| LV | carnivore | *Gaurochromis hiatus* | Gahi_A_02 | 484 |
| LV | carnivore | *Gaurochromis hiatus* | Gahi_A_03 | 485 |
| LV | carnivore | *Haplochromis thereuterion* | Hath_A_01 | 165 |
| LV | carnivore | *Haplochromis thereuterion* | Hath_A_01 | 166 |
| LV | carnivore | *Haplochromis thereuterion* | Hath_A_01 | 167 |
| LV | herbivore | *Neochromis omnicaeruleus* | Neom_A_01 | 260 |
| LV | herbivore | *Neochromis omnicaeruleus* | Neom_A_02 | 261 |
| LV | herbivore | *Neochromis omnicaeruleus* | Neom_A_03 | 262 |
| LM | herbivore | *Labeotropheus trewavasae "Red Top Thumbi West"* | Latr_A_01 | 447 |
| LM | herbivore | *Labeotropheus trewavasae "Red Top Thumbi West"* | Latr_A_02 | 448 |

| | | | | |
|---|---|---|---|---|
| LM | herbivore | *Labeotropheus trewavasae "Red Top Thumbi West"* | Latr_A_03 | 449 |
| LM | herbivore | *Tropheops tropheops "Makokola"* | Trtr_A_01 | 500 |
| LM | herbivore | *Tropheops tropheops "Makokola"* | Trtr_A_02 | 509 |
| LM | herbivore | *Tropheops tropheops "Makokola"* | Trtr_A_03 | 503 |
| LM | herbivore | *Petrotilapia. sp. "Thick bars"* | Peth_A_01 | 505 |
| LM | herbivore | *Petrotilapia. sp. "Thick bars"* | Peth_A_02 | 507 |
| LM | herbivore | *Petrotilapia. sp. "Thick bars"* | Peth_A_03 | 537 |
| LM | herbivore | *Petrotilapia. sp. "Yellow chin Chewere"* | Peye_A_01 | 371 |
| LM | herbivore | *Petrotilapia. sp. "Yellow chin Chewere"* | Peye_A_02 | 373 |
| LM | herbivore | *Petrotilapia. sp. "Yellow chin Chewere"* | Peye_A_03 | 374 |
| LM | carnivore | *Sciaenochromis fryeri* | Scfr_A_01 | 154 |
| LM | carnivore | *Sciaenochromis fryeri* | Scfr_A_02 | 155 |
| LM | carnivore | *Sciaenochromis fryeri* | Scfr_A_03 | 156 |
| LM | carnivore | *Aulonocara hansbaenschi* | Auha_A_01 | 490 |
| LM | carnivore | *Aulonocara hansbaenschi* | Auha_A_02 | 491 |
| LM | carnivore | *Aulonocara hansbaenschi* | Auha_A_03 | 492 |
